# Transcriptome and evolutionary analysis of *Pseudotrichomonas keilini*, a free-living anaerobic eukaryote

**DOI:** 10.1101/2024.09.26.614816

**Authors:** Hend Abu-Elmakarem, Stephen J. Taerum, Celine Petitjean, Michael Kotyk, Christopher Kay, Ivan Čepička, David Bass, Gillian H. Gile, Tom A. Williams

## Abstract

The early evolution of eukaryotes and their adaptations to low-oxygen environments are fascinating open questions in biology. Genome-scale data from novel eukaryotes, and particularly from free-living lineages, are key to answering these questions. The Parabasalia are an ancient lineage of anaerobes, and the most speciose lineage of Metamonada, a major lineage of eukaryotes. The most well-studied metamonads are parasitic parabasalids including *Trichomonas vaginalis, Tritrichomonas foetus*, and *Giardia intestinalis*, but very little genome-scale data is available for free-living members of the group. Here, we sequenced the transcriptome of *Pseudotrichomonas keilini*, a free-living parabasalian. Comparative genomic analysis indicated that *P. keilini* possesses a metabolism and gene complement that are in many respects similar to its parasitic relative *Trichomonas vaginalis*, and that in the time since their most recent common ancestor, it is the *Trichomonas vaginalis* lineage that has experienced more genomic change, likely due to the transition to a parasitic lifestyle. Features shared between *P. keilini* and *Trichomonas vaginalis* include a hydrogenosome (anaerobic mitochondrial homologue) that we predict to function much as in *Trichomonas vaginalis*, and a complete glycolytic pathway that is likely to represent one of the primary means by which *P. keilini* obtains ATP. Phylogenomic analysis indicates that *P. keilini* branches within a clade of endobiotic parabasalids, consistent with the hypothesis that different parabasalid lineages evolved towards parasitic or free-living lifestyles from an endobiotic, anaerobic or microaerophilic common ancestor.

## Introduction

Animals, plants and fungi are well-studied by biologists and genomes for many lineages are now available. However, most eukaryotic diversity is microbial, and many groups are poorly sampled by genomics and transcriptomics (Sibbald and Archibald, 2017). Among unicellular lineages, parasites are best represented by sequencing efforts. These include the parabasalid *Trichomonas vaginalis*, the causative agent of the most common non-viral sexually transmitted disease, trichomoniasis (Donné, 1836, WHO, 2023). *Trichomonas vaginalis* infects the genitourinary tracts of 187 million people every year around the world (Menezes et al., 2016) and raises the transmission of human immunodeficiency virus HIV (Petrin et al., 1998). There is therefore great interest in the biology of *Trichomonas vaginalis* and also in the evolution of parasitism in the whole Parabasalia.

There are over 450 described species of parabasalids. The majority are endobiotic, including parasites such as *Trichomonas vaginalis* and its relatives that infect other animals (Adl et al., 2007; Brugerolle and Lee, 2000; Cepicka et al., 2010; Yamin and MA, 1979). The first free-living parabasalid to be discovered is *Pseudotrichomonas keilini* (Bishop, 1939, 1935). *P. keilini* was originally isolated from pond water in Lincolnshire, UK, and later isolated from mangrove sediments in Japan, a lake in Cyprus (Yubuki et al., 2010), from fresh water sediments in Azerbaijan, sulfurous freshwater spring and brackish sediments in Greece, and from inland salt March in Spain (Céza et al., 2022). As parabasalids are thought to be ancestrally endobiotic (Čepička et al., 2017), several questions naturally arise about a free-living member of the group: did the free-living lifestyle of *P. keilini* evolve secondarily, or are transitions between free-living and host-associated lifestyles more common than anticipated in parabasalids? To answer these questions, we carried out transcriptome sequencing and bioinformatics analyses.

Here, we present a largely complete transcriptome dataset of *P. keilini*, a free-living anaerobic parabasalid isolated from salt marsh sediment in Spain. The transcriptome of *P. keilini* codes for 18,851 unique proteins. The annotation of these proteins shows that *P. keilini* has metabolic capabilities that are highly similar to its parasitic relative *Trichomonas vaginalis*. Based on analysis of gene content, *P. keilini* possesses a hydrogenosome used for energy production. As expected, our analyses suggest that carbohydrates provide the main source of energy to produce pyruvate via glycolysis which then enters the hydrogenosome to produce ATP, CO_2_ and H_2_. We envisage that the *P. keilini* transcriptome will be of use in further analyses of early eukaryote genomic and metabolic evolution.

## Methods

### Strain isolation and culture

*Pseudotrichomonas keilini* strain TOLEDOT was isolated from salt marsh sediment at Toledo, Castile-La Mancha, Spain, coordinates 39°58’42’’N, 3°39’20’’W. It was maintained as a mono-eukaryotic, polyxenic culture in Dobell-Laidlaw biphasic (Dobell and Laidlaw, 1926) (Dobell and Laidlaw 1926) medium at room temperature and subcultured every two weeks. Living and protargol-stained cells of *P. keilini* were examined under a microscope 1200 BX51 (Olympus) equipped with an Olympus DP71 camera, using DIC optics for living cells. Protargol-stained preparations were prepared as follows: moist films spread on coverslips were prepared from pelleted cultures obtained by centrifugation at 500 g for 8 minutes. The films were fixed in Bouin-Hollande’s fluid for approximately 15 hours, washed with 70% ethanol, and stained with 1% protargol (Bayer, I. G. Farbenindustrie) following (Nie, 1950) protocol. Micrographs of protargol-stained cells exhibiting the morphological characteristics of *P. keilini* are given in Supplementary Figure 1.

### Transcriptome sequencing and assembly

For transcriptome sequencing, RNA was extracted from 8 mL *P. keilini* culture using Trizol reagent (Invitrogen) and cDNA was synthesised using the SmartSeq2 protocol (Picelli et al. 2014) using poly-dT primers for first strand synthesis to enrich for eukaryotic transcripts and a template-switching oligo to enable sequencing of the 5’ end of each transcript. Libraries were built at the ASU genomics core using KAPA Biosystem’s LTP library preparation kit (KK8232). Library fragment size was analysed by Tapestation (Agilent), and quantified by qPCR (KAPA Library Quantification Kit, KK4835) on Thermo Fisher Scientific’s Quantstudio 5 before sequencing on an Illumina MiSeq V2 using 2×250 paired end chemistry. Additional transcriptome sequencing from another 8 mL *P. keilini* culture was carried out at Génome Québec using the NEB stranded mRNA library protocol (New England Biolabs) for cDNA synthesis and library preparation before sequencing 50 million reads on the NovaSeq6000 platform using 2×100 chemistry. Transcript reads were assembled using Trinity RNA-Seq v2.8.4 (Haas et al., 2013). Predicted proteins that were 100% identical over the overlapping length were clustered using CD-HIT 4.8.1 (Li and Godzik, 2006). Reads have been deposited in the NCBI Short Read Archive under the accession PRJNA884676.

### Gene finding and annotation

Protein prediction from transcripts was done using TransDecoder version 5.5.0 (Haas et al., 2013). Using default parameters, we obtained a total of 83,266 proteins from the total set of assembled transcripts. As we were unable to obtain a pure culture of *P. keilini*, we applied a conservative BLAST-based filter to these proteins to distinguish bona fide *P. keilini* proteins from prokaryotic contaminants of the assembly. To do so, we searched the predicted proteins against a custom database containing protein sets from 38 published genomes of excavates, along with a representative sampling of 73 other eukaryotes, 148 bacteria and 146 archaea (see Supplementary Table S1). Proteins for which either (i) the best hit was from a parabasalid relative of *P. keilini*, or (ii) alternatively where the first three database hits were eukaryotic in a Diamond BLAST search were retained as putative *P. keilini* proteins. We assigned functions to the filtered *P. keilini* protein set using eggNOG-mapper v2, which uses precomputed protein families from the eggNOG database (Huerta-Cepas et al., 2019, 2017, 2016) to perform functional annotation. For proteins which could not be annotated with eggNOG, we used blastKOALA to predict functions from the KEGG database (Kanehisa et al., 2017, 2016; Kanehisa and Goto, 2000).

### Phylogenetics

The species tree was inferred using SpeciesRax (Morel et al., 2022), based on 13,346 gene family trees clustered from 43 eukaryotic genomes using Broccoli v1.2 (Derelle et al., 2020). The branch lengths represent substitutions per site. The individual gene family trees were inferred using IQ-TREE2 (Minh et al., 2020), with 1000 rapid bootstraps, and the best-fitting model in each case (including LG+C20 and LG+C60 as potential site-heterogeneous options), was chosen by the Bayesian Information Criterion. The phylogenetic birth-death model was fit in Count following the procedure suggested in the user manual (Csűös, 2010); the final model included three categories to model variation in branch-specific lengths and duplication rates across the species tree.

### Hydrogenosome detection

We used the supplementary table in (Stairs et al., 2015) which included key enzymes of each of the five classes of mitochondria and its related organelles classified by Muller et al. (2012). Using Diamond (Buchfink et al., 2015), we blasted the sequences against the database of *P. keilini* filtered proteins to identify key enzymes. Other protein sequences were added for these enzymes by doing a blast search against the NCBI nr database and selecting the top five hits from eukaryotes, alphaproteobacteria, and other bacteria. The retrieved sequences were then aligned using MAFFT v7.390 (Katoh and Standley, 2013). After checking the alignment, and selecting the most complete sequences, we then ran BMGE-1.12 (Criscuolo and Gribaldo, 2010) which trims the multiple sequence alignment to remove poorly aligned regions. The trimmed alignments were used to construct maximum-likelihood trees with IQ-TREE-v1.6.10 using the best model selected by ModelFinder (Kalyaanamoorthy et al., 2017; Nguyen et al., 2015). Finally, we used HMMER 3.2.1 (http://hmmer.org/) to search the filtered *P. keilini* transcriptome for homologues of mitochondrial carrier family transport proteins, with an E-value threshold of < 10^−5^. We used the PF00153 HMM profile for the MCF proteins which contains 125,808 sequences from 1,117 species including *Trichomonas vaginalis*. HMMER profiles are probabilistic profile hidden Markov models (Durbin et al., 1998; Eddy, 1998; Krogh et al., 1994) which are used to identify distant protein homologs.

## Results and Discussion

### *Transcriptome of* P. keilini

Transcriptome reads were assembled into 137,389 transcripts using Trinity RNA-Seq (Haas et al., 2013), and predicted proteins were obtained using TransDecoder. We performed BLAST-based filtration to remove contaminants from the assembly (see methods), and clustered identical overlapping proteins to obtain 18,851 non-redundant predicted proteins, of which functions could be predicted for 12,811.

To evaluate the completeness of the *P. keilini* transcriptome, we conducted a BUSCO analysis (Simão et al., 2015) using a set of 255 conserved eukaryotic genes. We recovered 20 single-copy complete BUSCOs in *P. keilini*, in comparison to 22 in *Trichomonas vaginalis*. Thus, we concluded that we produced a near-complete transcriptome which is likely to be informative about the evolution and metabolism of *P. keilini*.

### P. keilini *branches within a clade of parasitic parabasalids*

To investigate the relationship of *P. keilini* to other parabasalids and metamonads, we used SpeciesRax (Morel et al. 2022) to infer a species tree including *P. keilini*, its closest parabasalid relatives, and a representative sample of other eukaryotic lineages, using 13,346 gene families clustered from 43 genomes (Figure 1). Interestingly, the phylogeny indicates that *P. keilini* branches within a clade of parasitic parabasalids with very good support, as the sister lineage to a clade comprising *Trichomonas vaginalis, Trichomonas gallinae* and *Trichomonas tenax* (extended quadripartition internode certainty (EQPIC) 0.53, reflecting good agreement between this species tree node and the underlying quartets of the input gene family trees). The cattle and feline parasite *Tritrichomonas foetus* branches sister to this *P. keilini-Trichomonas vaginalis* clade (Figure 1). The phylogeny implies that *P. keilini* evolved a free-living lifestyle from a parasitic ancestor, or alternatively that there have at least two transitions to parasitism within this clade: once in the ancestor of *Tritrichomonas foetus* and once in the ancestor of the *Trichomonas* clade. However, our phylogeny lacks many free-living and commensal endobiotic Parabasalia due to the lack of molecular data. In 18S rRNA phylogenies, *Pseudotrichomonas* and *Lacusteria* are the two deepest branches within the order Trichomonadida. The position of Trichomonadida within Parabasalia is not resolved, but it might be the sister clade of Honigbergiellida, which contains several free-living species. The endobiotic *Trichomitus batrachorum* branches distantly and belongs to yet another order, the Hypotrichomonadida (Céza et al. 2022). Parasitic and free-living species seem to have arisen multiple times each within the predominantly commensal endobiotic Parabasalia.

**Figure 1:**
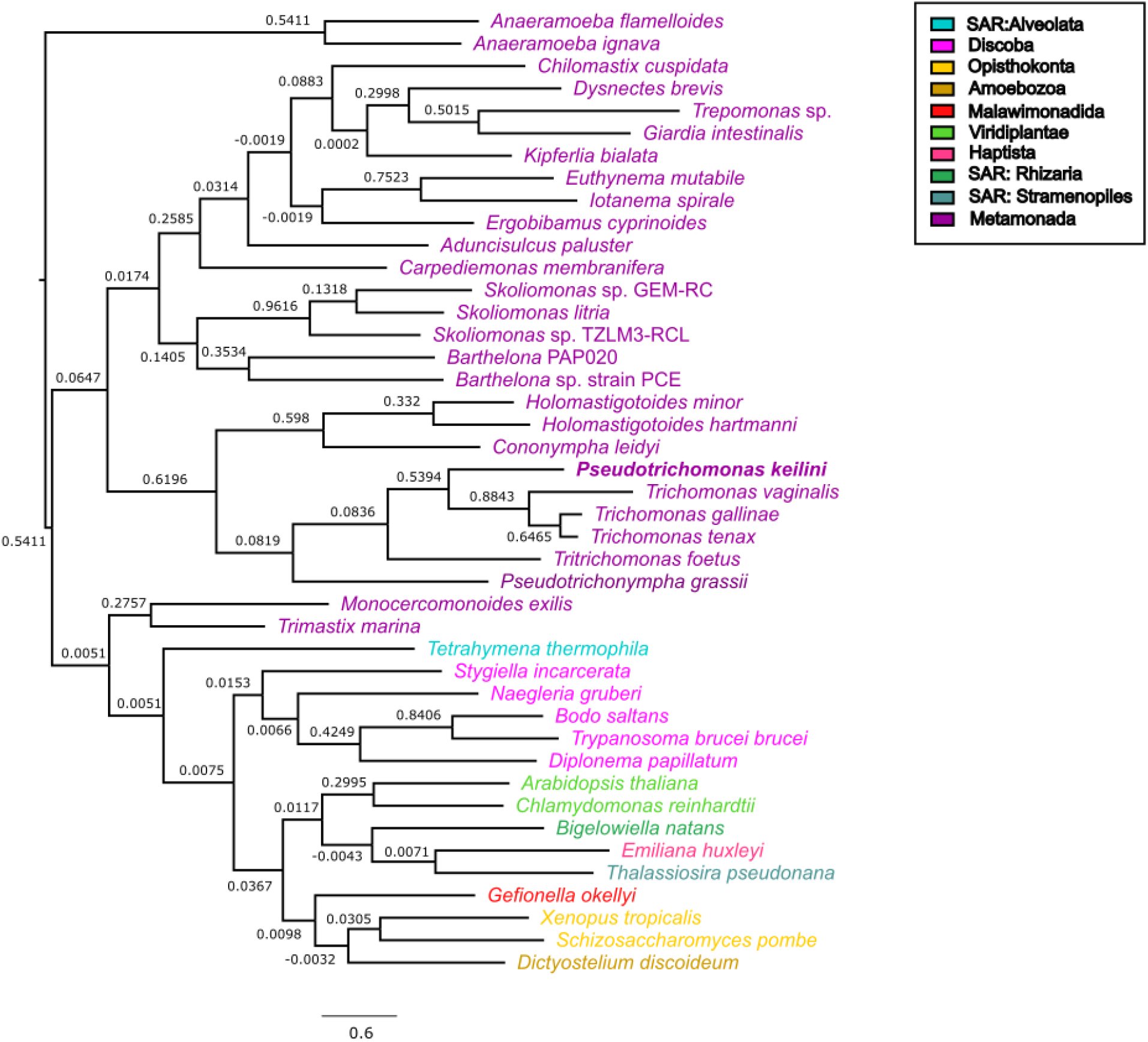
Species tree of *Pseudotrichomonas keilini* among parabasalids and metamonads. SpeciesRax estimated species tree from 13,346 gene family trees, clustered from 43 genomes). Branch supports are EQPIC scores (Morel et al. (2022), reflecting the degree of agreement between quartets in the input gene trees and the inferred maximum likelihood species tree; >0.5 denotes high support). *P. keilini* groups robustly with the parasitic parabasalids *Trichomonas vaginalis, Trichomonas foetus, Trichomonas gallinae*, and *Trichomonas tenax*. Branch lengths in units of mean expected substitutions per site. Species are coloured based on their taxonomic groups as indicated in the colour code box on the right.

More broadly, the maximum likelihood SpeciesRax topology recovered the monophyly of Metamonada (including Parabasalia, Fornicata, Preaxostyla, *Barthelona, Skoliomonas*, and Anaeramoebae), although the deep relationships between these lineages were poorly resolved, with EQPIC scores close to 0 for the deepest splits within Metamonada, representing conflicting support from gene family trees. Alternative approaches such as concatenation (Stairs et al., 2021; Williamson et al., 2024) or reconciliation methods that model gene tree uncertainty (Cerón-Romero et al., 2022; Morel et al., 2024) may be required to resolve these deep branches of the eukaryotic tree.

### *Gene content evolution in* P. keilini *and its relatives*

To investigate the evolutionary origins of the *P. keilini* genome, we mapped gene family evolution on the inferred eukaryotic species tree using a phylogenetic birth-death model implemented in Count (Csűös, 2010), summarised in Figure 2. The analysis suggested that the common ancestor of Parabasalia and Fornicata had a relatively small genome (2,975 gene families), with extensive gene gain in the parabasalid lineage after its divergence from the common ancestor with Fornicata. This is consistent with previous reports of gene family expansions in Parabasalia (Handrich et al., 2019; Maciejowski et al., 2023; Oyhenart and Breccia, 2014). The common ancestor of *P. keilini* and *Trichomonas vaginalis* was inferred to have a gene repertoire comparable in size to that of *P. keilini* (5,280 gene families), with additional gene gains in the *Trichomonas vaginalis* lineage after its divergence from *Trichomonas gallinae* and *Trichomonas tenax*. Functional annotation of genes gained in the *Trichomonas vaginalis* lineage highlighted a role for binding to cellular components of the urogenital tract, a crucial step for extracellular parasite survival (Pereira-Neves and Benchimol, 2007). Another key parasitic function can be seen through the presence of genes binding to spectrin, a protein found on the host cell surface which acts as the main gateway for target cell degradation (Fiori et al., 1997). These results are consistent with the view that the *Trichomonas vaginalis* lineage adapted to a parasitic lifestyle after divergence from its common ancestor with *P. keilini*, in part through the gene gains mapped here.

**Figure 2:**
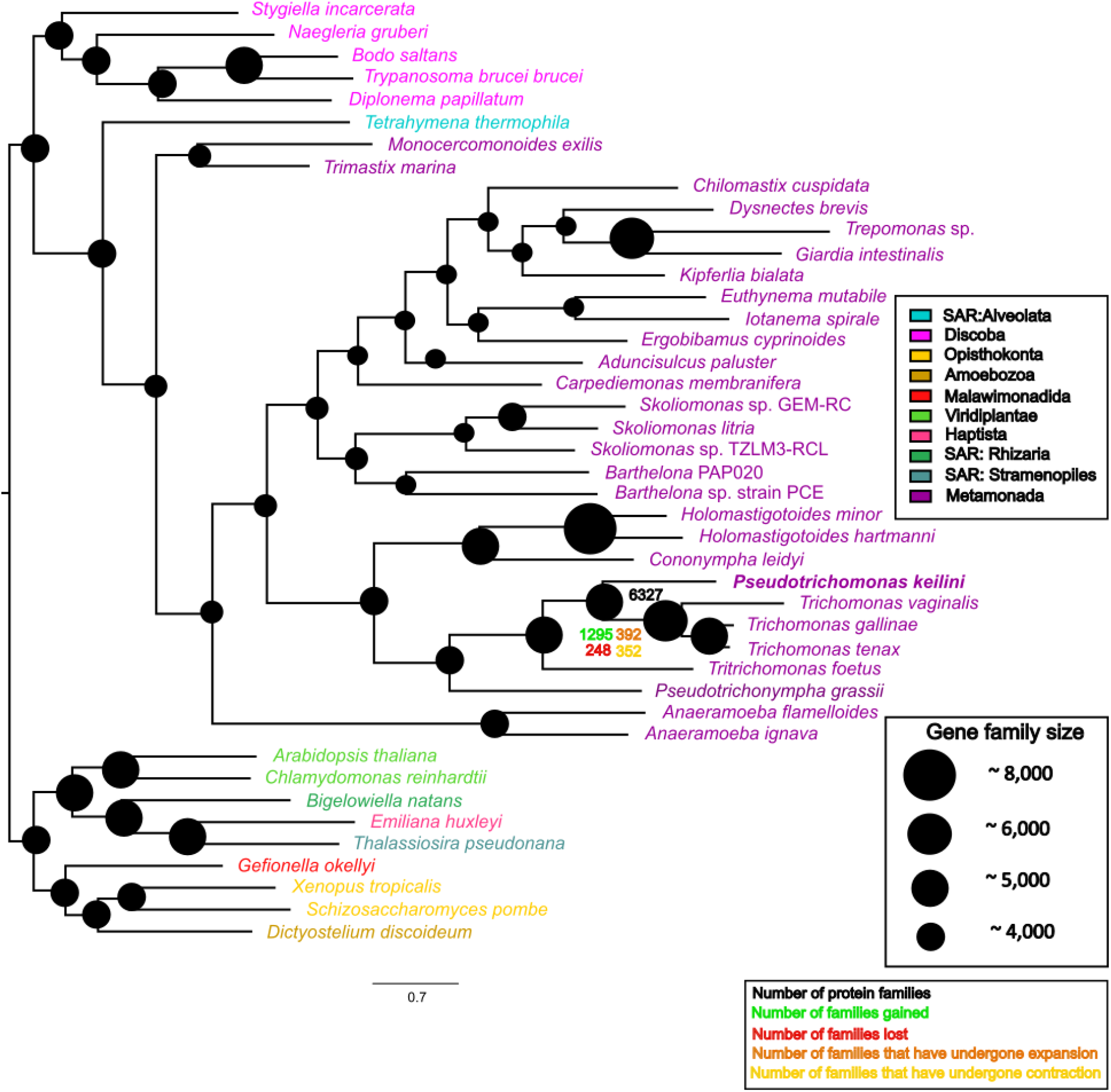
Gene family evolution in parabasalids and metamonads. We used a phylogenetic birth-death model implemented in Count (Csuros 2010) to map gene family evolution onto the inferred species tree. Numbers and the diameter of circles indicate gene family repertoire size at ancestral nodes, while family gains, losses, expansions and contractions are plotted for the *Trichomonas* lineage after its divergence from *P. keilini*. The analysis was also performed on a species tree manually edited to reflect the consensus view of deep eukaryotic relationships (Figure S2), with closely similar results for gene content evolution within metamonads.

As the maximum likelihood species tree contained some heterodox features, we also performed a sensitivity analysis in which we fit the phylogenetic birth-death model to a species tree edited to more closely reflect current views of eukaryotic phylogeny (reviewed in (Burki et al., 2020)), including a sister group relationship between parabasalids and Anaeramoebae, the grouping of *Tetrahymena thermophila* with other SAR members, and a specific relationship between Discoba and Metamonada); see Supplementary Figure 2 for the alternative tree. The results were closely similar.

### *The hydrogenosome of* P. keilini

Metamonads are ancestrally anaerobic, and characterised metamonads have a range of reduced mitochondrial homologues or mitochondria-related organelles (MROs; Stairs et al. 2015). One of the best-characterised MROs is the hydrogenosome of *Trichomonas vaginalis*, and so we sought to investigate the MRO of *P. keilini* and to compare it with that of its close anaerobic, parasitic relative. To do so, we searched the *P. keilini* protein set for proteins previously implicated in MRO function and metabolism (Stairs et al., 2015), proteins that have been localised to the hydrogenosome in *Trichomonas vaginalis* (569 proteins, Schneider et al., 2011) and other hallmark mitochondrial proteins including the mitochondrial carrier family of transporters.

This analysis confirmed that *P. keilini* possesses a hydrogenosome that, in many respects, is similar to that of *Trichomonas vaginalis*: both organisms encode the same subset of 22 hallmark proteins drawn from a larger set found in a wide range of MROs across the eukaryotic tree of life (Stairs et al. 2015), including the key enzymes needed to reduce protons to molecular hydrogen via the oxidative decarboxylation of pyruvate (pyruvate-ferredoxin oxidoreductase, [FeFe]-hydrogenase and the associated maturases) (Figure S4, Supp Tables S2,S3,S5). Overall, our analyses suggest that *P. keilini* has all the enzymes needed to carry out glycolysis, with the resulting pyruvate imported into the hydrogenosome for further catabolism.

The transition from mitochondrion to hydrogenosome was previously inferred to have occurred in the common ancestor of all metamonads (Leger et al., 2017). This is consistent with our findings, in that phylogenetic analysis of the key hydrogenosomal enzymes recovered a monophyletic clade of parabasalids in each of the trees (Supplementary Figures 3-14). In total, the *P. keilini* transcriptome encodes orthologues of 487/569 *Trichomonas vaginalis* proteins localised to the hydrogenosome by proteomics (Schneider et al., 2011). Both organisms also encode the same complement of six paralogous mitochondrial carrier family proteins, suggesting that the requirements for transport into and out of the MRO are similar in each case (see Supplementary Material for more discussion on hydrogenosomal import in *P. keilini*).

**Figure 3:**
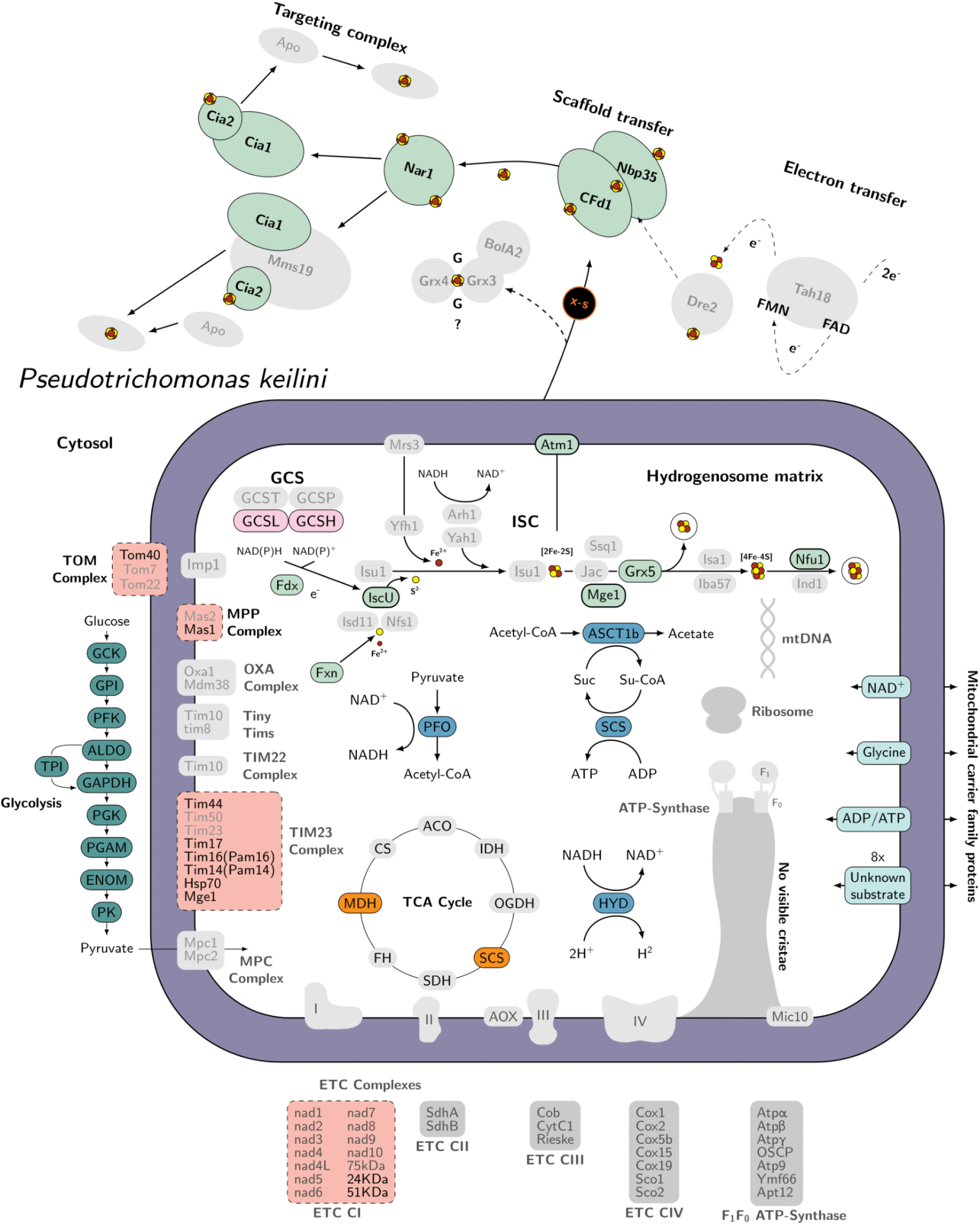
Predicted metabolic pathways in the *Pseudotrichomonas keilini* hydrogenosome. Key enzymatic reactions are depicted, including glycolysis, the main pathway used to produce pyruvate, which is then oxidised by the TCA cycle in the hydrogenosome. The translocases of the outer and inner mitochondrial membranes (TOM and TIM, respectively) and the ETC subunits that were detected in complex I are also indicated; note that the subunits identified (NuoE and NuoF) are common in anaerobes, and are involved in other processes beyond oxidative phosphorylation (Hrdy et al., 2004; Stairs et al., 2021). Boxes with solid outlines represent complexes for which all subunits were identified, dashed outlines represent complexes with some subunits identified, and boxes in grey with no outline indicate complexes with no subunits identified. Figure design is adapted from Peña-Diaz and Lukeš (2018) and Lewis et al. (2019) (Lewis et al., 2019; Peña-Diaz and Lukeš, 2018).

The biosynthesis of iron-sulphur (Fe-S) clusters is perhaps the most widely (although not universally) conserved function of mitochondria and mitochondria-related organelles (Stairs et al., 2015). The *P. keilini* hydrogenosome carries out Fe-S cluster biosynthesis using the ISC pathway, as in *Trichomonas vaginalis*. We detected orthologues of all the ISC enzymes present in *Trichomonas vaginalis* and *Naegleria gruberi* in the *P. keilini* transcriptome. Interestingly, *P. keilini* shares an iron–sulfur flavoprotein (Isf) of bacterial origin used in the detoxification of reactive oxygen species with *Trichomonas vaginalis* and one other eukaryotic anaerobe, *Entamoeba histolytica* (Peña-Diaz and Lukeš, 2018).

## Conclusion

Here, we sequenced the transcriptome of a free-living anaerobic parabasalid, *Pseudotrichomas keilini*. Although free-living, the gene content, metabolism and hydrogenosome of *P. keilini* are in many ways similar to its parasitic relative, *Trichomonas vaginalis*. The phylogenetic position of *P. keilini* in the species tree inferred from a large sample of protein-coding genes is consistent with the numerous transitions between free-living and host-associated lifestyles that have been suggested by 18S rRNA gene trees. Based on analysis of BUSCO gene content, the transcriptome is likely to be largely complete, and will represent a useful resource for future comparative analyses of parabasalids and eukaryotic evolution more broadly.

## Acknowledgments

We thank František Šťáhlavský for providing the salt marsh sediment from which this strain of *P. keilini* was isolated and Pierre Lafont for help with figure design. IČ and MK were supported by the Czech Science Foundation project no. 22-22538S. This work was supported in part by a grant from the US National Science foundation, DEB-2045329, to GG.

## Data availability

Read data has been deposited in the NCBI SRA (accession number PRJNA884676). The transcriptome assembly, gene models and annotations are available in a FigShare repository: https://doi.org/10.6084/m9.figshare.26528119.v1.

