## Supplementary Figures and Tables for "Transcriptome and evolutionary analysis of *Pseudotrichomonas keilini*, a free-living anaerobic eukaryote"

Supplementary Material for “Transcriptome and evolutionary analysis of *Pseudotrichomonas keilini,* a free-living anaerobic eukaryote*”.*

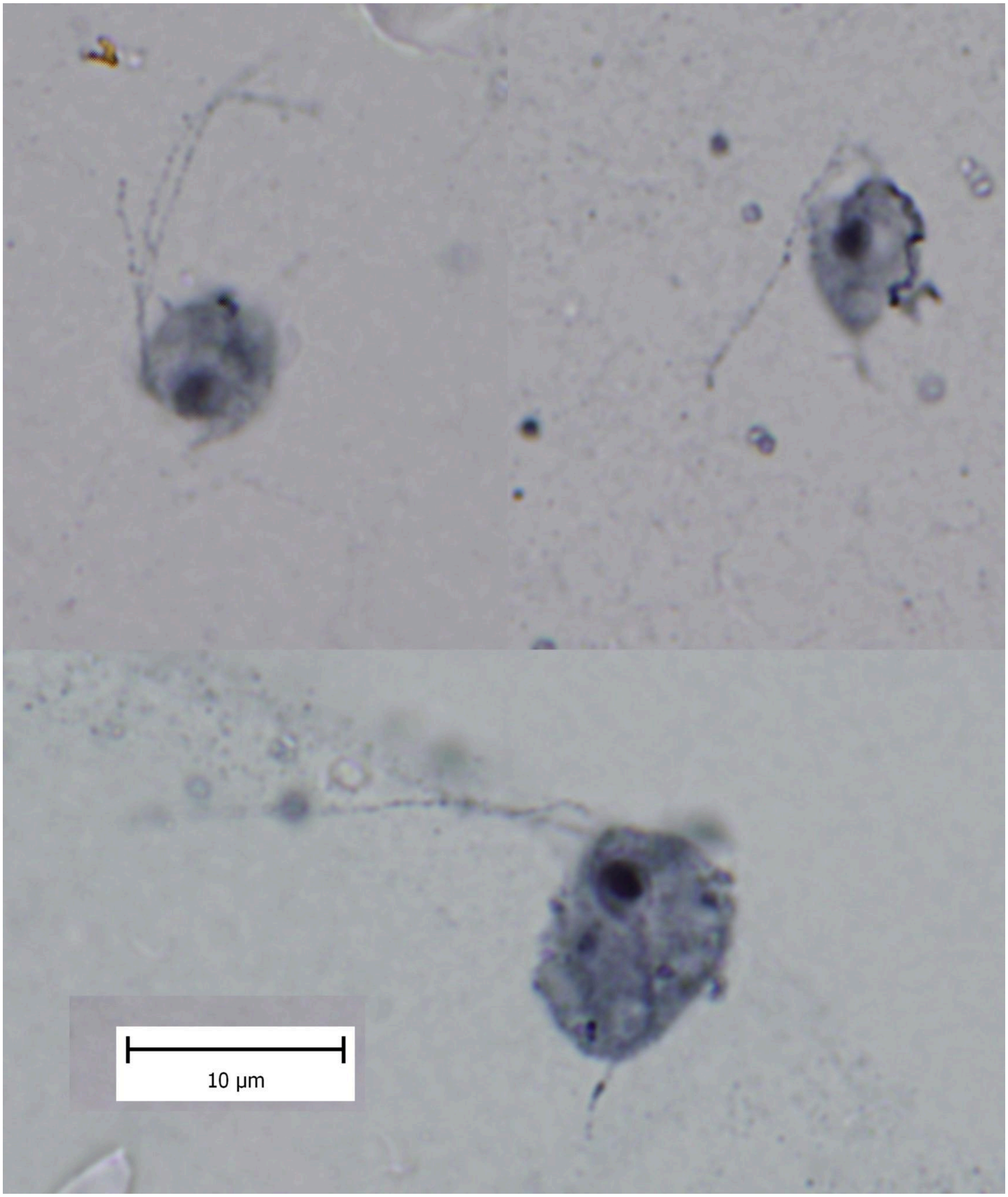

**Figure S1. Morphology of *Pseudotrichomonas keilini*.** Protargol-stained cells exhibit the diagnostic characters of this species: three anterior flagella (upper left cell), a well developed undulating membrane that does not continue as a free flagellum (upper right cell), a large parabasal body (this differentiates the organism from *Lacusteria*; lower cell), and a normally developed axostyle (lower cell).

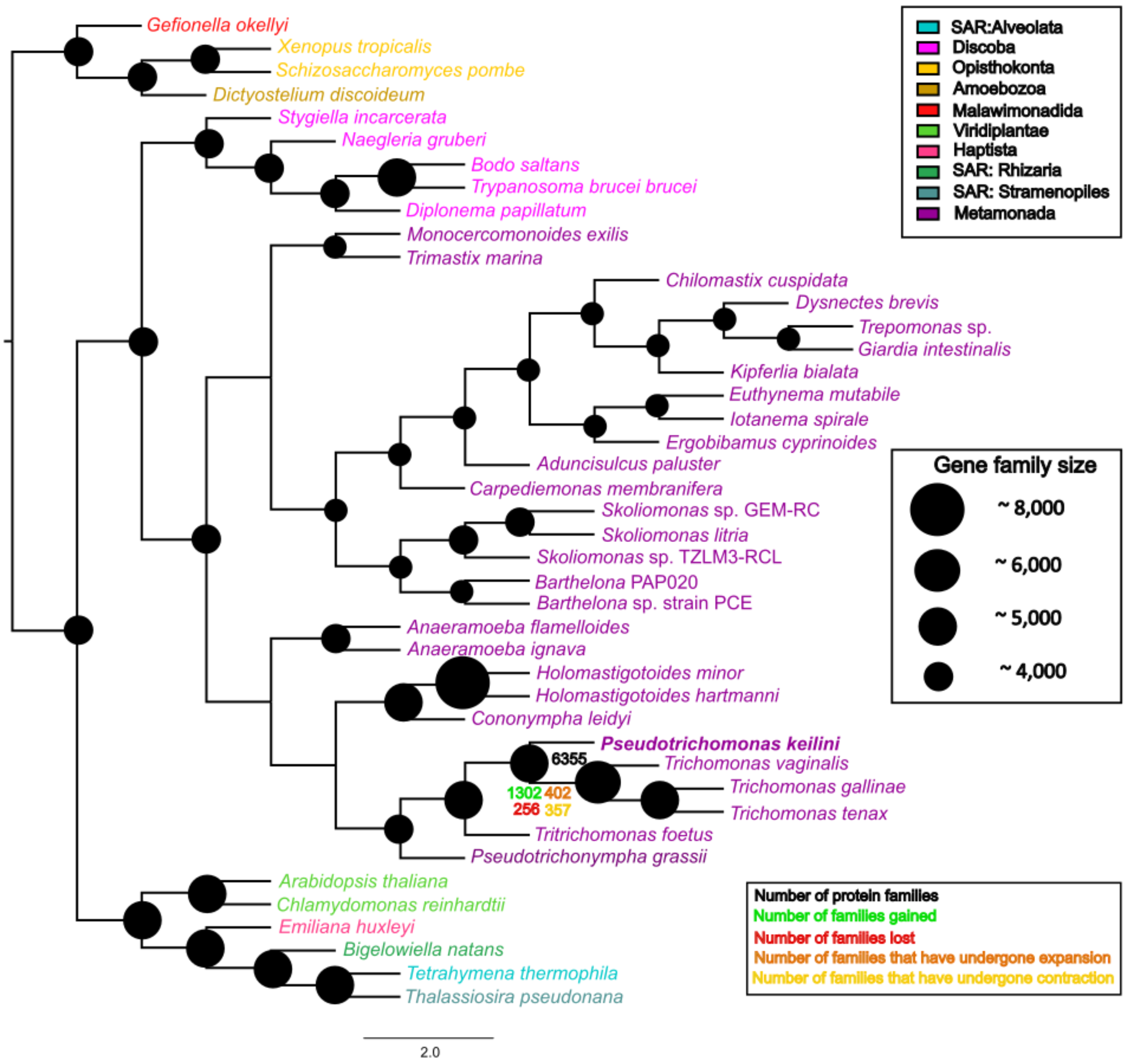

**Figure S2: Gene family evolution in parabasalids and metamonads.** We used a phylogenetic birth-death model implemented in Count (Csűös 2010) to map gene family evolution onto a species tree manually edited to reflect the consensus view of deep eukaryotic relationships. Numbers, and the diameter of circles, indicate gene family repertoire size at ancestral nodes, while family gains, losses, expansions and contractions are plotted for the *Trichomonas* lineage after its divergence from *P. keilini*.

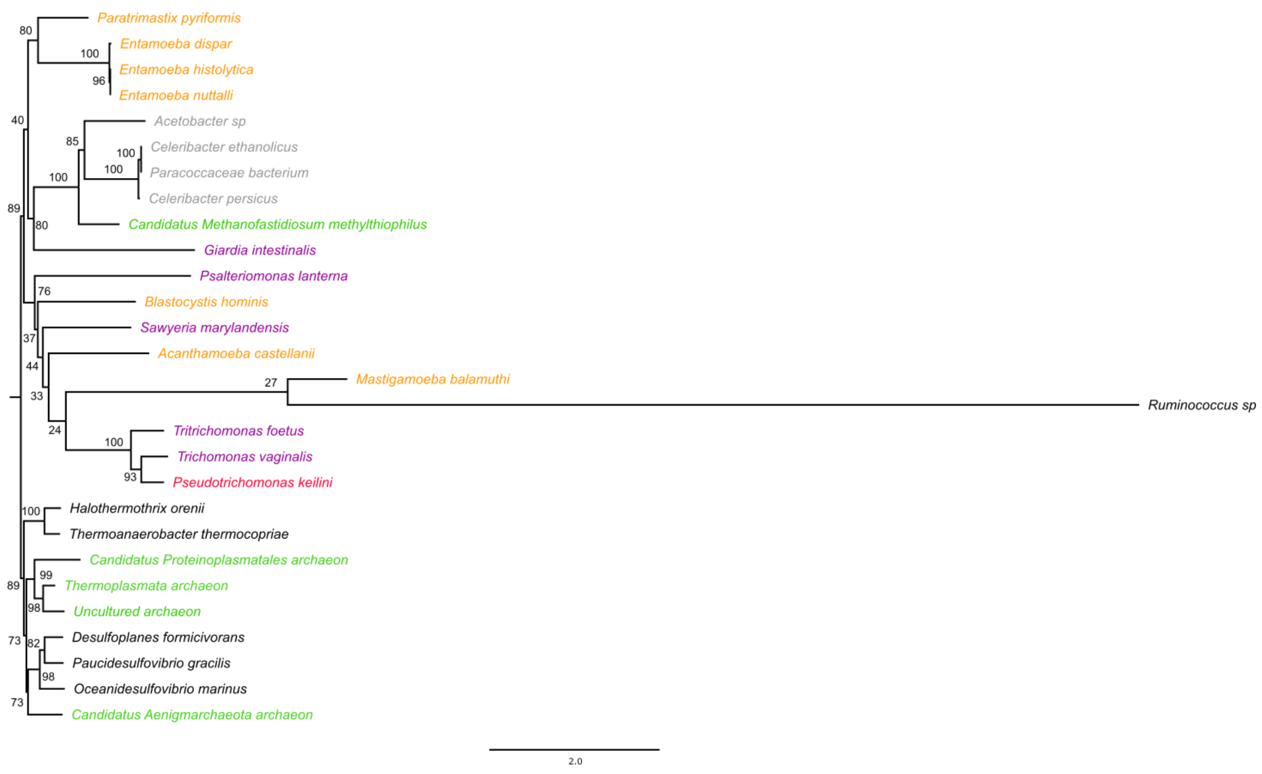

**Figure S3: Phylogeny of Pyruvate:ferredoxin oxidoreductase (PFO).** PFO is a key hydrogenosomal enzyme that catalyzes the interconversion of pyruvate to Acetyl-CoA. Most sampled excavate sequences including *Psalteriomonas lanterna* and *Sawyeria marylandensis* form a clade with the parabasalids *P. keilini, T. vaginalis,*and *T. foetus.* The maximum likelihood gene tree was inferred using IQ-TREE. Branch supports are ultrafast bootstrap values, and branch lengths are proportional to the expected number of substitutions per site, as indicated by the scale bar.

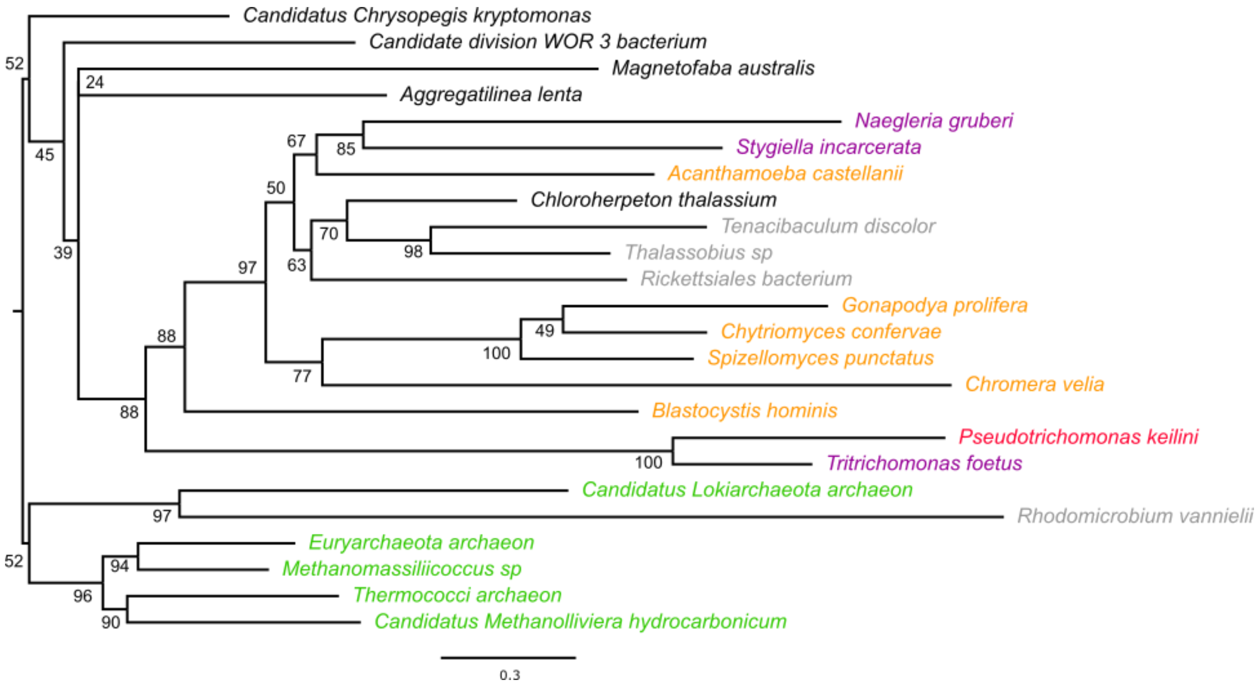

**Figure S4: Phylogenetic analysis of Acetyl:succinate CoA-transferase subunit b (ASCT1b)** which is a bidirectional enzyme that acts on the conversion of succinate to acetyl-CoA and vice versa. In the tree, *P. keilini* groups with *T. foetus* and they both form part of a larger clade which includes other eukaryotes, some of which are excavates. The maximum likelihood gene tree was inferred using IQ-TREE. Branch supports are ultrafast bootstrap values, and branch lengths are proportional to the expected number of substitutions per site, as indicated by the scale bar.

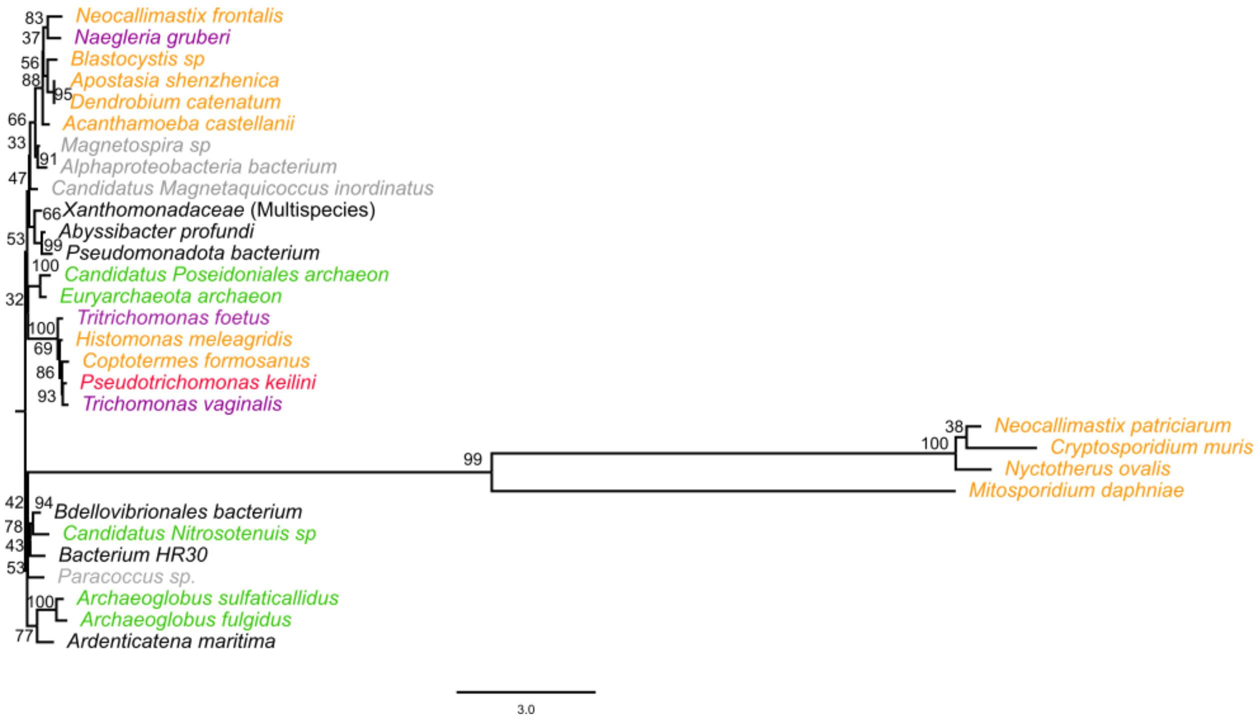

**Figure S5: Phylogenetic analysis of Succinyl coenzyme A synthetase (SCS)** which is a Krebs cycle enzyme that catalyzes the interconversion of succinyl-CoA to succinate. *T. foetus, T. vaginalis*, and *P. keilini*, but not the eukaryotes as a whole, form a clade. The maximum likelihood gene tree was inferred using IQ-TREE. Branch supports are ultrafast bootstrap values, and branch lengths are proportional to the expected number of substitutions per site, as indicated by the scale bar.

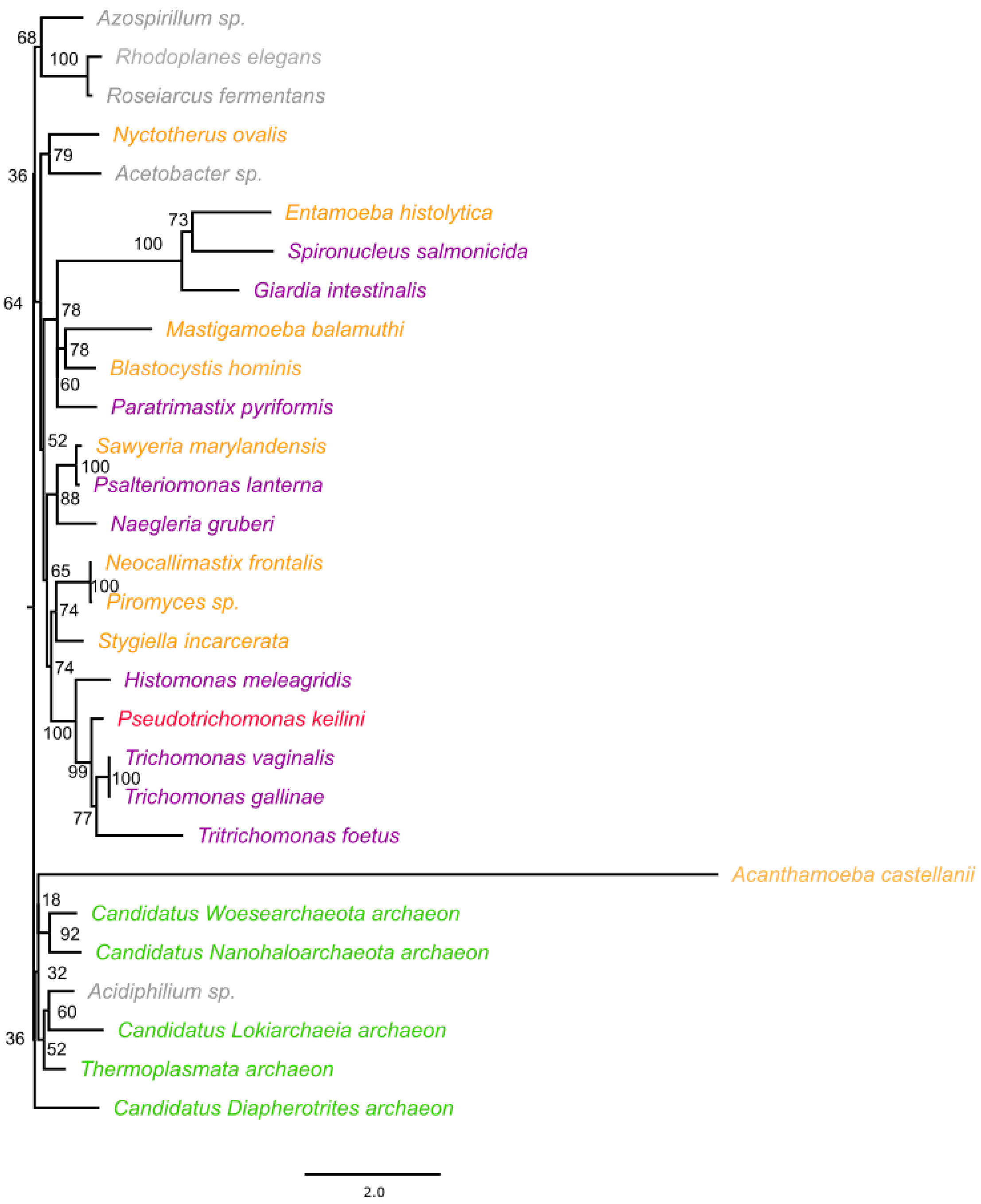

**Figure S6: Phylogeny of Fe-Fe hydrogenase (hydA) enzyme** which is responsible for the production of molecular hydrogen and one of the key hydrogenosomal enzmes. Another sequence from the excavate *Histomonas meleagridi* groups with the parabasalian clade of *P. keilini, T. vaginalis,* and *T. foetus, Trichomonas sp* and *T. gallinae*. The maximum likelihood gene tree was inferred using IQ-TREE. Branch supports are ultrafast bootstrap values, and branch lengths are proportional to the expected number of substitutions per site, as indicated by the scale bar.

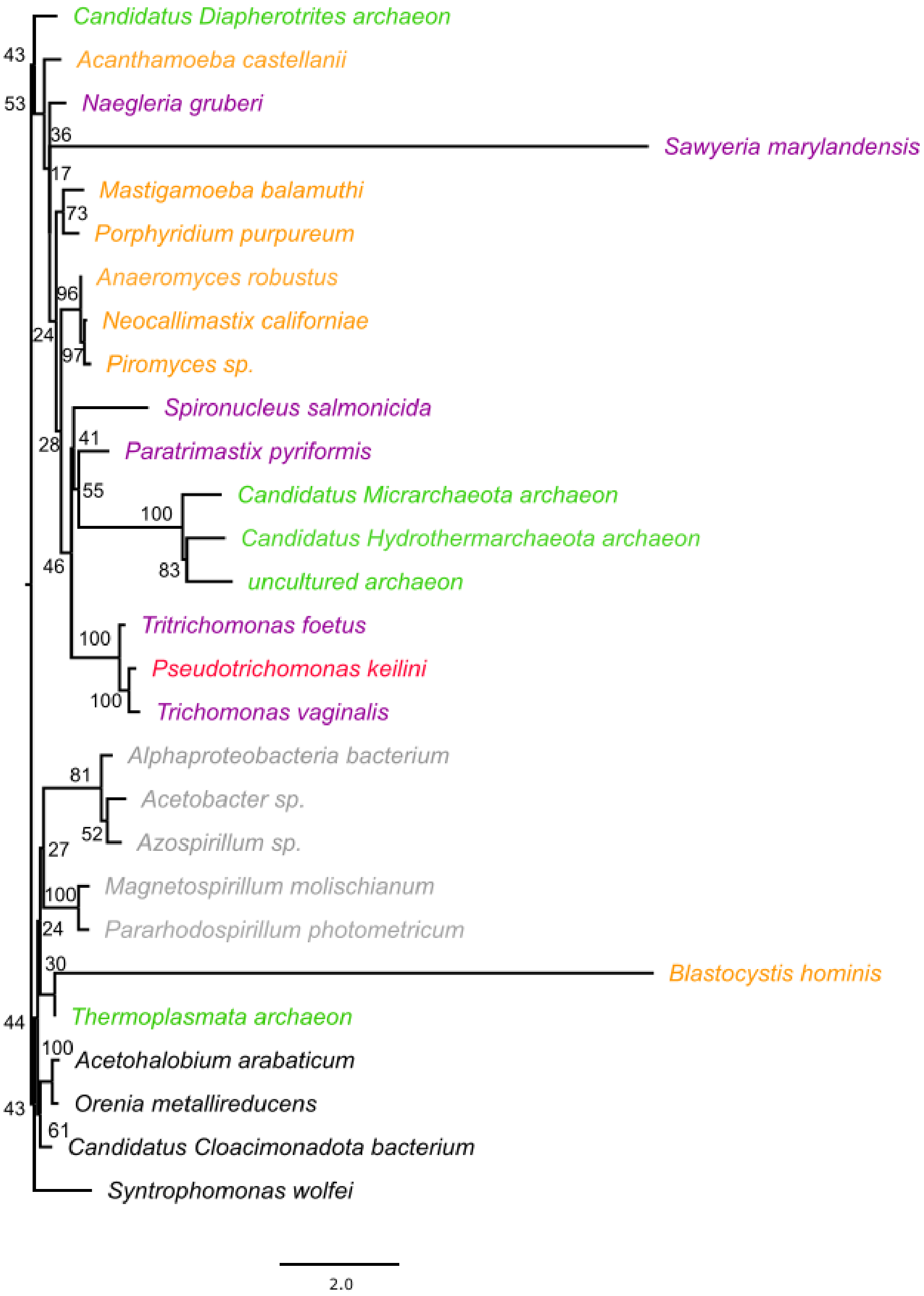

**Figure S7: Phylogeny of radical SAM domain containing protein (hydE)**, one of the three maturase enzymes required for the synthesis of a mature Fe-Fe-hydrogenase(hydA). In this tree, the three parabasalids of *P. keilini, T. vaginalis,* and *T. foetus* group together. The maximum likelihood gene tree was inferred using IQ-TREE. Branch supports are ultrafast bootstrap values, and branch lengths are proportional to the expected number of substitutions per site, as indicated by the scale bar.

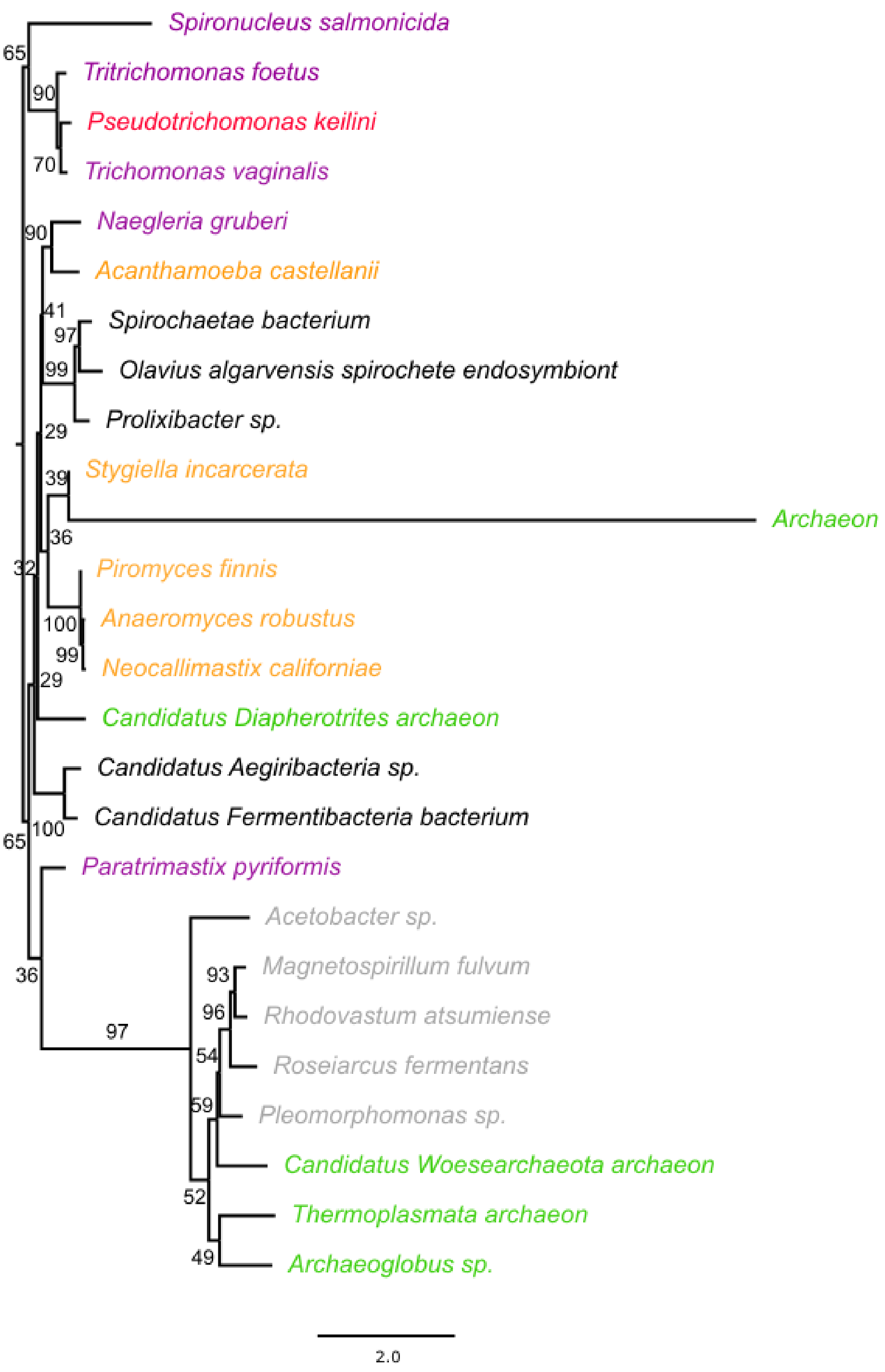

**Figure S8: Phylogeny of small GTP-binding protein (hydF)**, another one of the three maturase enzymes required for the synthesis of a mature Fe-Fe-hydrogenase(hydA). In this tree, the three parabasalids of *P. keilini, T. vaginalis*, and *T. foetus* group together. The maximum likelihood gene tree was inferred using IQ-TREE. Branch supports are ultrafast bootstrap values, and branch lengths are proportional to the expected number of substitutions per site, as indicated by the scale bar.

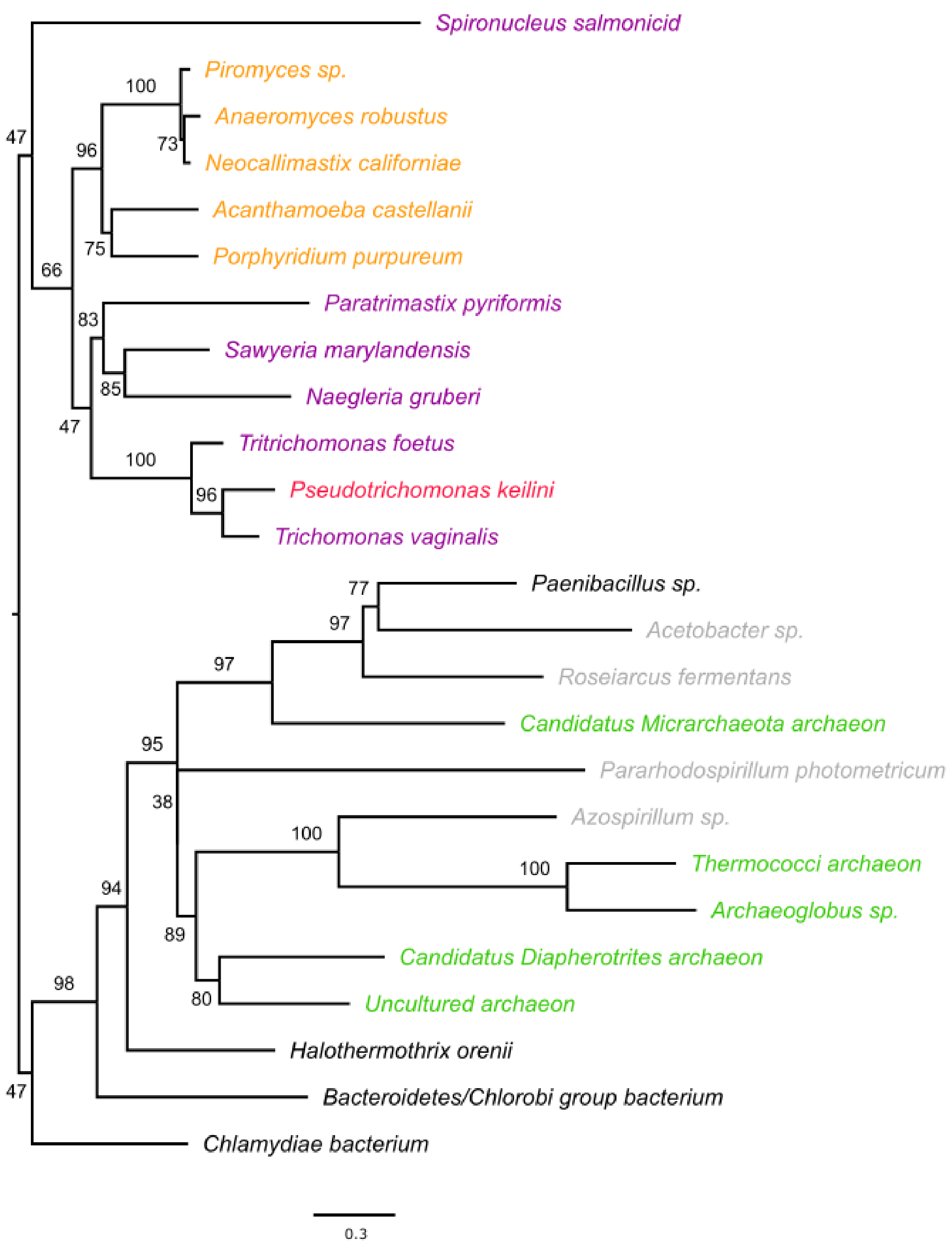

**Figure S9: Phylogeny of FeFe-hydrogenase assembly protein (hydG),** the third maturase enzyme required for the synthesis of a mature Fe-Fe-hydrogenase(hydA). A clade of the three parabasalids *P. keilini, T. vaginalis*, and *T. foetus* can be seen which is part of a bigger clade of eukaryotes including excavates. The maximum likelihood gene tree was inferred using IQ-TREE. Branch supports are ultrafast bootstrap values, and branch lengths are proportional to the expected number of substitutions per site, as indicated by the scale bar.

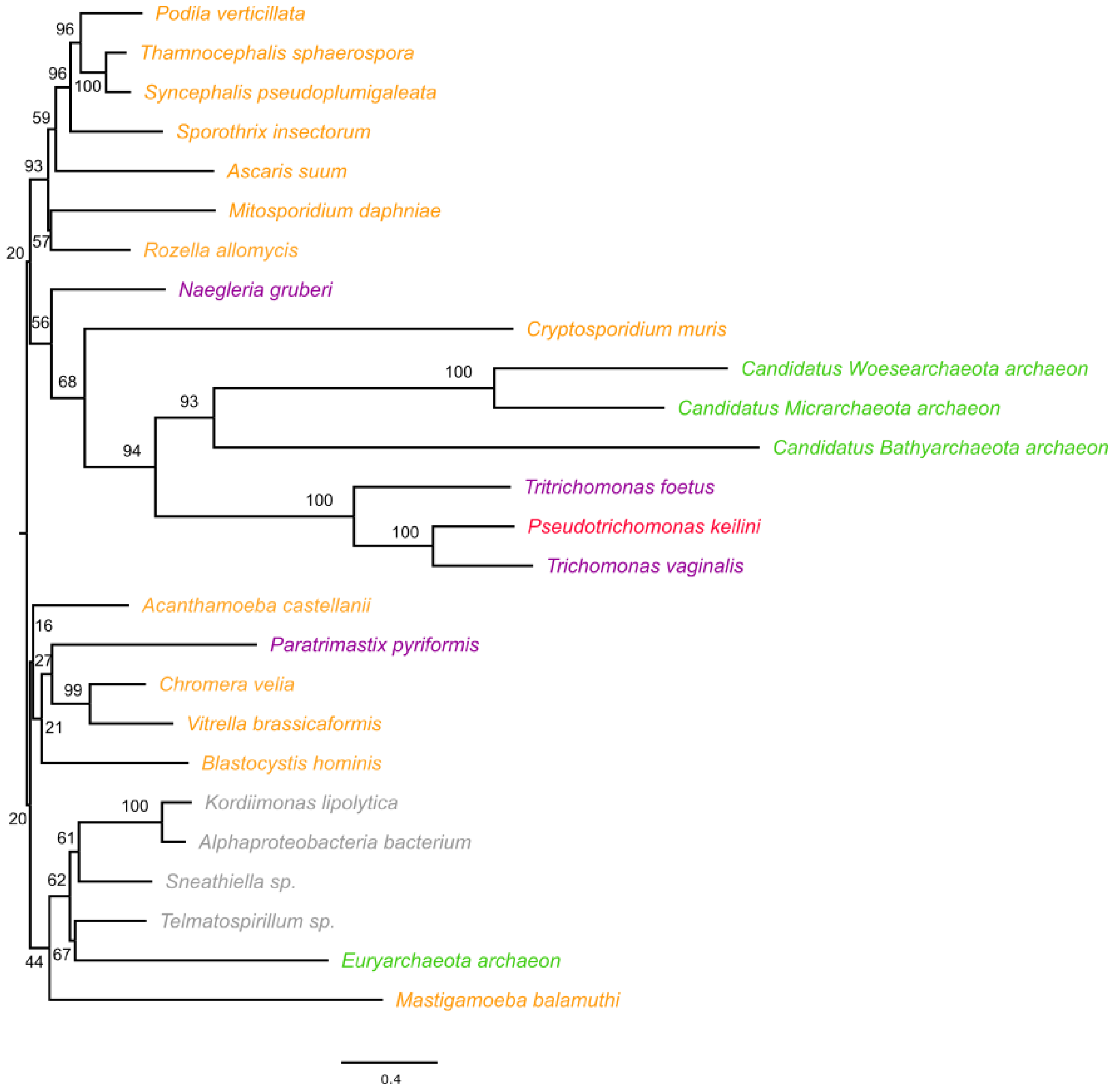

**Figure S10: Phylogeny of L-protein (GCSL),** which is part of the glycine cleavage system. The tree topology shows a clade of parabaslids consisting of *P. keilini, T. vaginalis*, and *T. foetus*, while other eukaryotic sequences form a separate clade. The maximum likelihood gene tree was inferred using IQ-TREE. Branch supports are ultrafast bootstrap values, and branch lengths are proportional to the expected number of substitutions per site, as indicated by the scale bar.

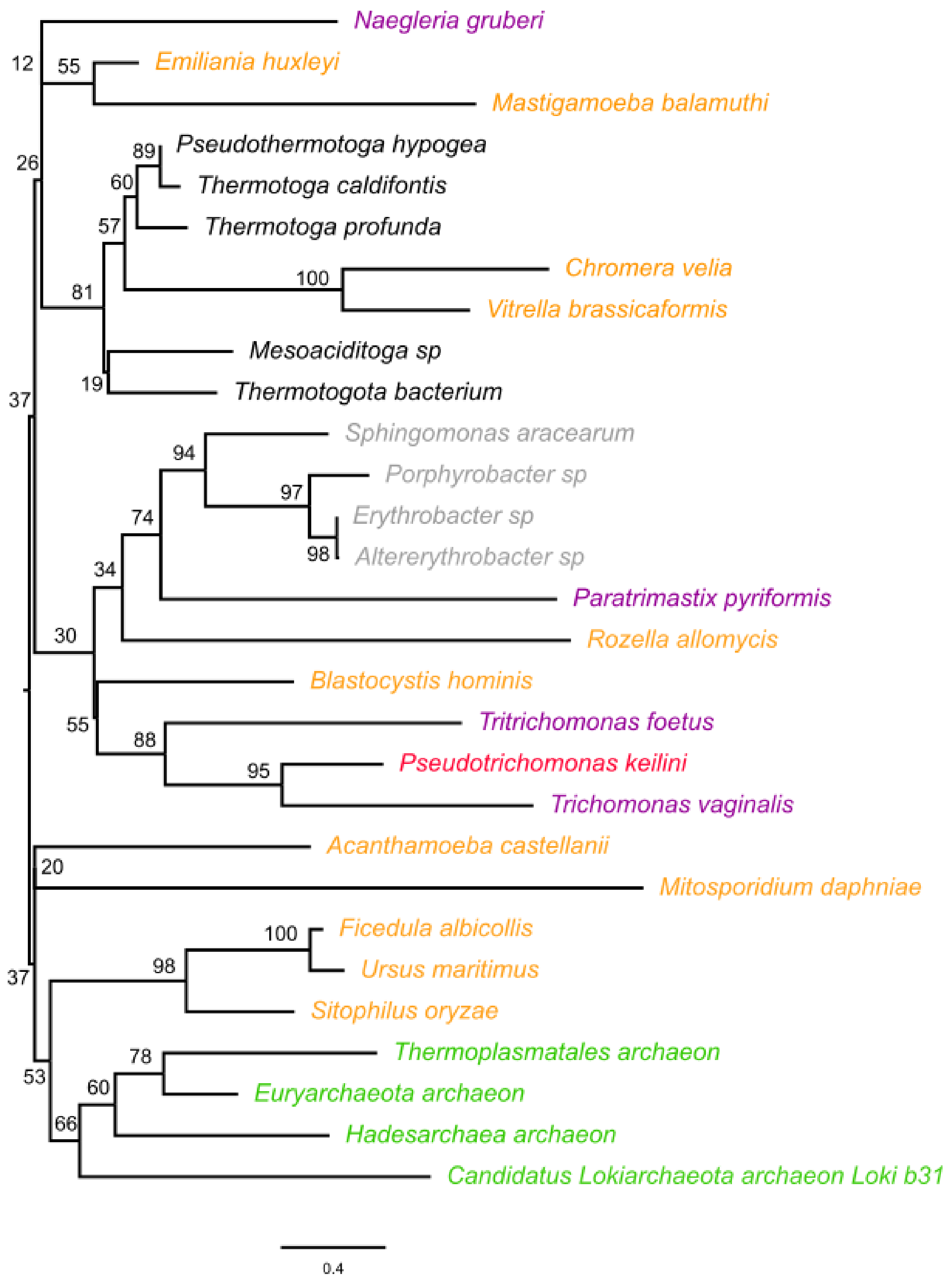

**Figure S11: Phylogeny of H-protein (GCSH)**, the second enzyme detected in *P. keilini*'s glycine cleavage system. The tree contains a clade of parabaslids including *P. keilini, T. vaginalis,* and *T. foetus* that is part of a larger clade comprising other eukaryotic and some alphaproteobacterial sequences, which could indicate an alphaproteobacterial origin of the enzyme. The maximum likelihood gene tree was inferred using IQ-TREE. Branch supports are ultrafast bootstrap values, and branch lengths are proportional to the expected number of substitutions per site, as indicated by the scale bar.

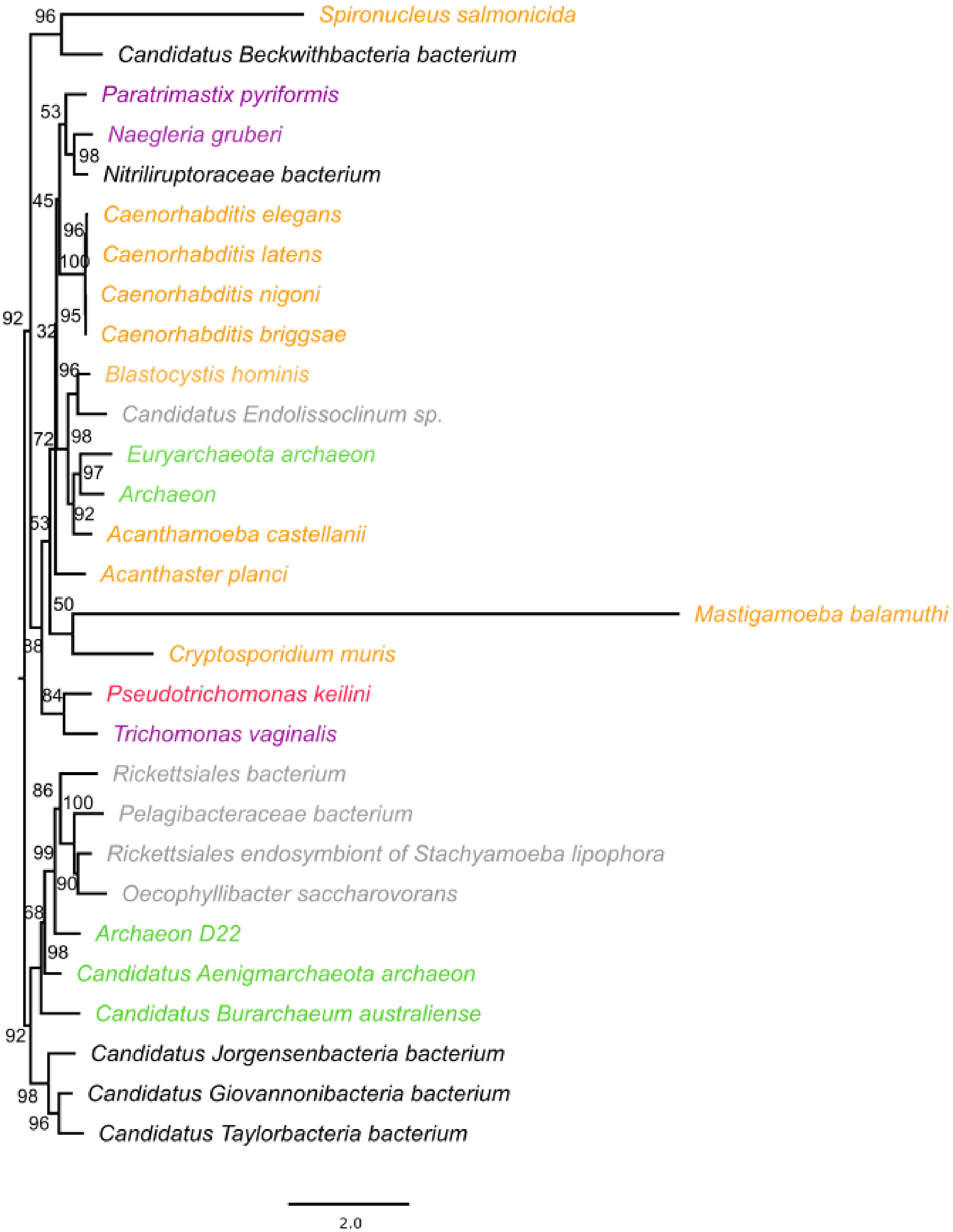

**Figure S12: Phylogenetic analysis of Serine hydroxymethyltransferase (SHMT) enzyme** which also plays a role in the glycine system. The tree shows that neither eukaryotes nor excavates are monophyletic. However, *P. Keilini* and *T. vaginalis* are grouping together. The maximum likelihood gene tree was inferred using IQ-TREE. Branch supports are ultrafast bootstrap values, and branch lengths are proportional to the expected number of substitutions per site, as indicated by the scale bar.

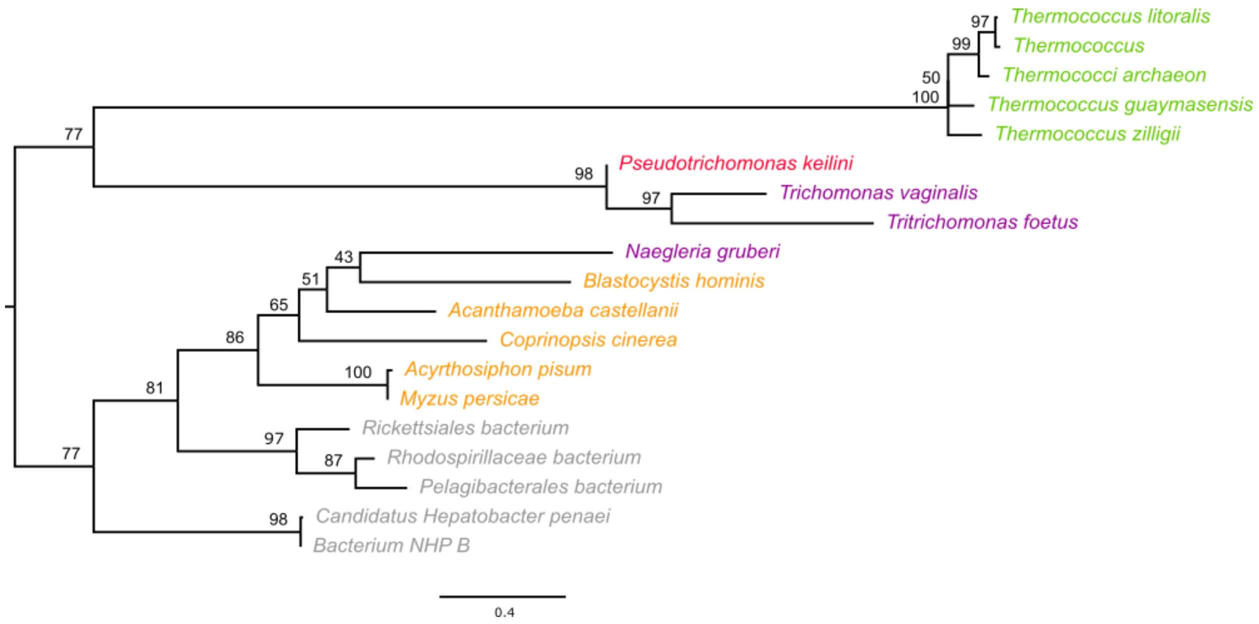

**Figure S13: Phylogeny of 24-kDa NADH-quinone oxidoreductase subunit E (NUOE),** one of the components of Complex I of the electron transport chain. In this tree, the *P. keilini* sequence is grouping with the other parabasalian ones of *T. vaginalis* and *T. foetus*, and not with alphaproteobacteria. The maximum likelihood gene tree was inferred using IQ-TREE. Branch supports are ultrafast bootstrap values, and branch lengths are proportional to the expected number of substitutions per site, as indicated by the scale bar.

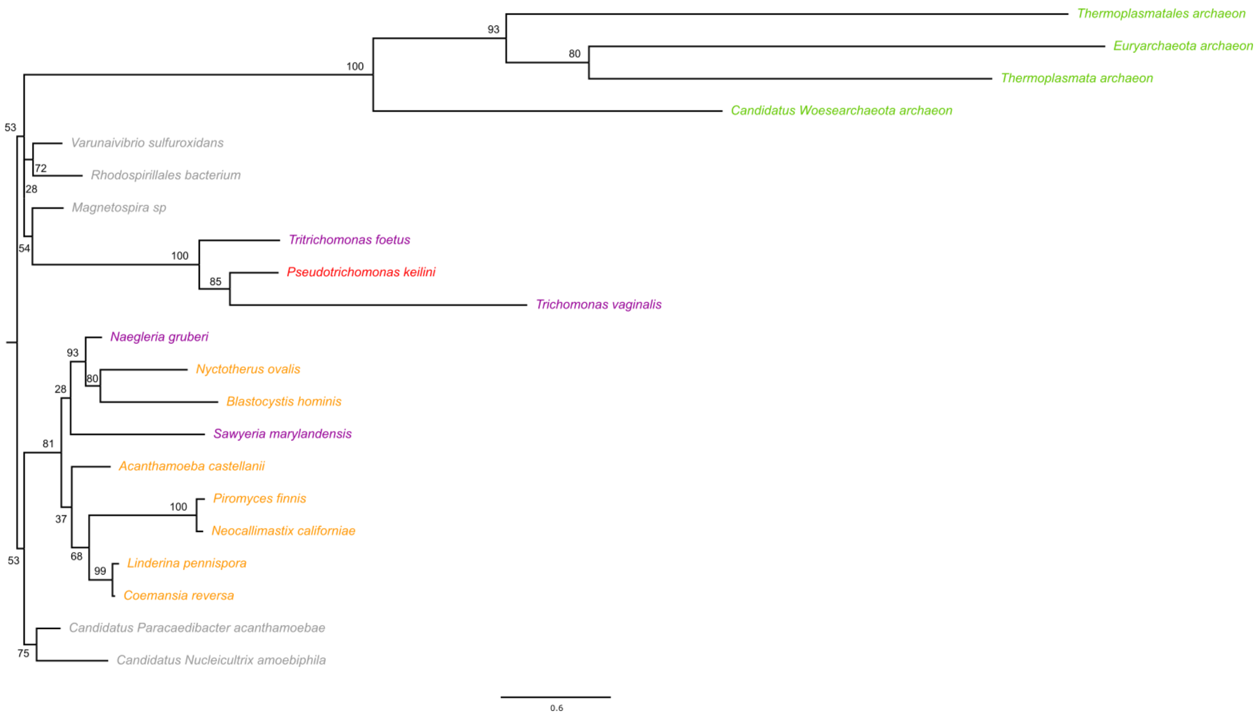

**Figure S14: Phylogenetic tree of 51-kDa NADH-quinone oxidoreductase subunit F (NUOF),** another component of Complex I of the electron transport chain. The tree topology shows that the parabasalian group of *P. keilini, T. vaginalis*, and *T. foetus* are grouping together within a larger clade of alphaproteobacterial sequences. This could indicate an endosymbiotic origin of the enzyme. The maximum likelihood gene tree was inferred using IQ-TREE. Branch supports are ultrafast bootstrap values, and branch lengths are proportional to the expected number of substitutions per site, as indicated by the scale bar.

Supplementary Table S1: List of species used in the filtration of eukaryotic proteins from contaminants

| Species name | Domain | Taxonomy |
|---|---|---|
| *Cyanophora paradoxa* | Eukaryotes | Archaeplastida |
| *Chondrus crispus* | | |
| *Galdieria sulphuraria* | | |
| *Cyanidioschyzon merolae* | | |
| *Porphyra umbilicalis* | | |
| *Chlamydomonas reinhardtii* | | |
| *Volvox carteri* | | |
| *Tetraselmis sp* | | |
| *Monoraphidium neglectum* | | |
| *Bathycoccus prasinos* | | |
| *Ostreococcus tauri* | | |
| *Micromonas pusilla* | | |
| *Auxenochlorella protothecoides* | | |
| *Oryza sativa* | | |
| *Arabidopsis thaliana* | | |
| *Physcomitrella patens* | | |
| *Selaginella moellendorffii* | | |
| *Klebsormidium nitens* | | |
| *Carpediemonas membranifera* | | Excavates |
| *Dysnectes brevis* | | |
| *Ergobibamus cyprinoides* | | |
| *Kipferlia bialata* | | |
| *Trepomonas sp* | | |
| *Aduncisulcus paluster* | | |
| *Chilomastix cuspidata* | | |
| *Trimastix marina* | | |
| *Naegleria gruberi* | | |
| *Stygiella incarcerata* | | |
| *Bodo saltans* | | |
| *Diplonema papillatum* | | |
| *Euglena gracilis* | | |
| *Giardia lamblia ATCC50803* | | |
| *Leishmania braziliensis* | | |

| | | |
|---|---|---|
| *Leishmania donovani species complex* | | |
| *Leishmania infantum JPCM5* | | |
| *Leishmania major Friedlin* | | |
| *Leishmania mexicana MHOM GT 2001 U1103* | | |
| *Leishmania panamensis* | | |
| *Leptomonas pyrrhocoris* | | |
| *Leptomonas seymouri* | | |
| *Phytomonas sp isolateEM1* | | |
| *Trypanosoma brucei brucei* | | |
| *Trypanosoma brucei gambiens* | | |
| *Trypanosoma congolense IL3000* | | |
| *Trypanosoma cruzi CLBrener* | | |
| *Trypanosoma cruzi Dm28c* | | |
| *Trypanosoma cruzi ID25* | | |
| *Trypanosoma cruzi marinkellei* | | |
| *Trypanosoma equiperdum* | | |
| *Trypanosoma grayi* | | |
| *Trypanosoma rangeli SC58* | | |
| *Trypanosoma theileri* | | |
| *Trypanosoma vivax Y486* | | |
| *Trichomonas vaginalis G3* | | |
| *Tritrichomonas foetus* | | |
| *Monocercomonoides* | | |
| *Ectocarpus siliculosus* | | **SAR** |
| *Thalassiosira pseudonana* | | |
| *Fragilariopsis cylindrus* | | |
| *Nannochloropsis gaditana* | | |
| *Thraustotheca clavata* | | |
| *Phaeodactylum tricornutum* | | |
| *Thalassiosira oceanica* | | |
| *Aureococcus anophagefferens* | | |
| *Leptocylindrus danicus* | | |
| *Odontella aurita* | | |
| *Nitzschia RCC80* | | |
| *Bolidomonas pacifica* | | |
| *Fistulifera solaris* | | |

| | | |
|---|---|---|
| *Tetrahymena thermophila* | | |
| *Paramecium tetraurelia* | | |
| *Stylonychia lemnae* | | |
| *Protoceratium reticulatum* | | |
| *Noctiluca scintillans* | | |
| *Togula jolla* | | |
| *Polarella glacialis* | | |
| *Amphidinium carterae* | | |
| *Oxytricha trifallax* | | |
| *Stentor coeruleus* | | |
| *Bigelowiella natans* | | |
| *Reticulomyxa filosa* | | |
| *Elphidium margaritaceum* | | |
| *Cryptophyceae sp* | | |
| *Guillardia theta* | | **Cryptista** |
| *Goniomonas avonlea* | | |
| *Emiliana huxleyi* | | |
| *Chrysochromulina sp* | | |
| *Raphidiophrys heterophryoidea* | | **Haptista** |
| *Acanthocystis sp* | | |
| *Choanocystis sp* | | |
| *Raineriophrys erinaceoides* | | |
| *Allomyces macrogynus* | | |
| *Schizosaccharomyces pombe* | | |
| *Spizellomyces punctatus* | | |
| *Lobosporangium transversale* | | |
| *Parvularia atlantis* | | |
| *Fonticula alba* | | |
| *Amphimedon queenslandica* | | **Obazoa** |
| *Limulus polyphemus* | | |
| *Xenopus tropicalis* | | |
| *Nematostella vectensis* | | |
| *Lottia gigantea* | | |
| *Monosiga brevicollis* | | |
| *Salpingoeca rosetta* | | |

| | | |
|---|---|---|
| *Gefionella okellyi* | | **Malawimonadidae** |
| *Ancoracysta twista* | | **Janouskovec** |
| *Ancyromonas sigmoides* | | **Eukaryota incertae sedis; Ancoracysta** |
| *Fabomonas tropica* | | **Ancyromonadida; Planomonadidae; Fabomonas** |
| *Nutomonas longa* | | **Ancyromonadida; Nutomonas** |
| *Diphylleia rotans* | | **CRuMs; Collodictyonidae; Diphylleia** |
| *Rigifila ramosa* | | **CRuMs; Rigifilida; Rigifila** |
| *Candidatus Berkelbacteria bacterium GWA2_46_7* | **Bacteria** | **CPR Berkelbacteria** |
| *Candidatus Berkelbacteria bacterium GWA2_35_9* | | **CPR Berkelbacteria** |
| *Candidatus Berkelbacteria bacterium GWA2_38_9* | | **CPR Berkelbacteria** |
| *Candidatus Berkelbacteria bacterium GWAE1_39_12* | | **CPR Berkelbacteria** |
| *Candidatus Berkelbacteria bacterium CG1_02_42_45* | | **CPR Berkelbacteria** |
| *Candidatus Berkelbacteria bacterium CG2_30_39_44* | | **CPR Berkelbacteria** |
| *Candidatus Berkelbacteria bacterium CG2_30_43_20* | | **CPR Berkelbacteria** |
| *Candidatus Berkelbacteria bacterium RIFCSPLOWO2_01_FULL_50_28* | | **CPR Berkelbacteria** |
| *Candidatus Berkelbacteria bacterium RIFOXYA2_FULL_43_10* | | **CPR Berkelbacteria** |

| | | |
|---|---|---|
| *Candidatus Candidate division WS6 bacterium GW2011_GWA2_37_6* | | **CPR Dojkabacteria** |
| *Candidate division WS6 bacterium GW2011_GWF1_35_23* | | **CPR Dojkabacteria** |
| *Candidate division Kazan bacterium GW2011_GWA1_50_15* | | **CPR Kazan** |
| *Candidate division Kazan bacterium GW2011_GWA1_44_22* | | **CPR Kazan** |
| *Candidatus Amesbacteria bacterium RIFOXYB1_FULL_44_23* | | **CPR Microgenomates** |
| *Candidatus Beckwithbacteria bacterium GW2011_GWC1_49_16* | | **CPR Microgenomates** |
| *Candidatus Collierbacteria bacterium RIFOXYB1_FULL_49_13* | | **CPR Microgenomates** |
| *Candidatus Curtissbacteria bacterium RBG_16_39_7* | | **CPR Microgenomates** |
| *Candidatus Daviesbacteria bacterium RIFCSPHIGHO2_12_FULL_37_16* | | **CPR Microgenomates** |
| *Candidatus Gottesmanbacteria bacterium GW2011_GWA1_43_11* | | **CPR Microgenomates** |
| *Microgenomates group bacterium GW2011_GWA2_46_16* | | **CPR Microgenomates** |
| *Candidatus Woykebacteria bacterium GWB1_45_5* | | **CPR Microgenomates** |
| *Candidatus Levybacteria bacterium RIFCSPHIGHO2_02_FULL_37_10* | | **CPR Microgenomates** |
| *Candidatus Pacebacteria bacterium CG1_02_43_31* | | **CPR Microgenomates** |
| *Candidatus Woykebacteria bacterium RBG_13_40_15* | | **CPR Microgenomates** |
| *Candidatus Woykebacteria bacterium RBG_13_40_7b* | | **CPR Microgenomates** |
| *Candidatus Roizmanbacteria bacterium RIFCSPLOWO2_01_FULL_45_11* | | **CPR Microgenomates** |
| *Candidatus Shapirobacteria bacterium GW2011_GWF2_37_20* | | **CPR Microgenomates** |

| | | |
|---|---|---|
| *Candidatus Woesebacteria bacterium GW2011_GWD2_40_19* | | **CPR Microgenomates** |
| *Candidatus Woesebacteria bacterium RIFCSPHIGHO2_01_FULL_41_10* | | **CPR Microgenomates** |
| *Candidatus Falkowbacteria bacterium GW2011_GWE2_38_254* | | **CPR Parcubacteria** |
| *Candidatus Giovannonibacteria bacterium RIFCSPHIGHO2_02_43_16* | | **CPR Parcubacteria** |
| *Candidatus Jorgensenbacteria bacterium GWA1_54_12* | | **CPR Parcubacteria** |
| *Candidatus Jorgensenbacteria bacterium RIFCSPHIGHO2_02_FULL_45_20* | | **CPR Parcubacteria** |
| *Candidatus Kaiserbacteria bacterium RIFCSPHIGHO2_01_FULL_56_24* | | **CPR Parcubacteria** |
| *Parcubacteria bacterium SCGC AAA011-A09* | | **CPR Parcubacteria** |
| *Candidatus Magasanikbacteria bacterium GW2011_GWA2_45_39* | | **CPR Parcubacteria** |
| *Candidatus Magasanikbacteria bacterium GW2011_GWA2_46_17* | | **CPR Parcubacteria** |
| *Candidatus Moranbacteria bacterium RIFCSPHIGHO2_01_FULL_55_24* | | **CPR Parcubacteria** |
| *Candidatus Moranbacteria bacterium GW2011_GWC2_37_8* | | **CPR Parcubacteria** |
| *Candidatus Nomurabacteria bacterium CG1_02_31_12* | | **CPR Parcubacteria** |
| *Candidatus Nomurabacteria bacterium RIFCSPLOWO2_01_FULL_36_10b* | | **CPR Parcubacteria** |
| *Candidatus Uhrbacteria bacterium RIFCSPLOWO2_02_FULL_49_11* | | **CPR Parcubacteria** |
| *Candidatus Uhrbacteria bacterium RIFCSPLOWO2_02_FULL_54_37* | | **CPR Parcubacteria** |

| | | |
|---|---|---|
| *Candidatus Uhrbacteria bacterium RIFOXYC2_FULL_47_19* | | **CPR Parcubacteria** |
| *Candidatus Wolfebacteria bacterium RIFOXYB1_FULL_54_12* | | **CPR Parcubacteria** |
| *Candidatus Yanofskybacteria bacterium RIFCSPHIGHO2_01_FULL_44_17* | | **CPR Parcubacteria** |
| *Candidatus Peregrinibacteria bacterium CG1_02_41_10* | | **CPR Peregrinibacteria** |
| *Candidatus Peregrinibacteria bacterium CG1_02_54_53* | | **CPR Peregrinibacteria** |
| *Candidatus Saccharibacteria bacterium CG2_30_41_52* | | **CPR Saccharibacteria** |
| *Candidatus Saccharibacteria bacterium RIFCSPHIGHO2_12_FULL_49_19* | | **CPR Saccharibacteria** |
| *Candidate division WWE3 bacterium RIFCSPLOWO2_01_FULL_42_11* | | **CPR WWE3** |
| *Chloroherpeton thalassium ATCC 35110* | | **FBC Chlorobi** |
| *Gemmatimonadetes bacterium RIFCSPLOWO2_12_FULL_68_9* | | **FBC Gemmatimonadetes** |
| *Ignavibacteria bacterium GWA2_55_25* | | **FBC Ignavibacteria** |
| *Candidate division Zixibacteria bacterium SM23_73_2* | | **FBC Zixibacteria** |
| *Thermithiobacillus tepidarius DSM 3134* | | **Proteobacteria Acidithiobacillia** |
| *Kordiimonas gwangyangensis DSM 19435* | | **Proteobacteria AlphaProteobacteria** |
| *Pelagibacterium halotolerans B2* | | **Proteobacteria AlphaProteobacteria** |
| *Rhodobacteraceae bacterium CG2_30_10_405* | | **Proteobacteria AlphaProteobacteria** |
| *Granulibacter bethesdensis CGDNIH1* | | **Proteobacteria AlphaProteobacteria** |

| | | |
|---|---|---|
| *Tistrella mobilis KA081020 65* | | **Proteobacteria AlphaProteobacteria** |
| *Anaplasma marginale* | | **Proteobacteria AlphaProteobacteria** |
| *Citromicrobium sp. JLT1363* | | **Proteobacteria AlphaProteobacteria** |
| *Bordetella bronchiseptica RB50* | | **Proteobacteria BetaProteobacteria** |
| *Comamonas testosteroni TK102* | | **Proteobacteria BetaProteobacteria** |
| *Gallionella capsiferriformans ES 2* | | **Proteobacteria BetaProteobacteria** |
| *Methylovorus sp. SIP3 4* | | **Proteobacteria BetaProteobacteria** |
| *Deefgea rivuli DSM 18356* | | **Proteobacteria BetaProteobacteria** |
| *Bacteriovorax marinus SJ* | | **Proteobacteria DeltaProteobacteria** |
| *Desulfotignum phosphitoxidans FiPS 3* | | **Proteobacteria DeltaProteobacteria** |
| *Myxococcus xanthus DK 1622* | | **Proteobacteria DeltaProteobacteria** |
| *Syntrophus aciditrophicus SB* | | **Proteobacteria DeltaProteobacteria** |
| *Lebetimonas sp. JS032* | | **Proteobacteria EpsilonProteobacteria** |
| *Anaerobiospirillum succiniciproducens DSM 6400* | | **Proteobacteria GammaProteobacteria** |
| *Haliea rubra CM41 15a DSM* | | **Proteobacteria GammaProteobacteria** |
| *Arhodomonas aquaeolei DSM 8974* | | **Proteobacteria GammaProteobacteria** |
| *Arsenophonus nasoniae DSM 15247* | | **Proteobacteria GammaProteobacteria** |
| *Legionella micdadei ATCC 33218* | | **Proteobacteria GammaProteobacteria** |
| *Methyloglobulus morosus KoM1* | | **Proteobacteria GammaProteobacteria** |

| | | |
|---|---|---|
| *Saccharospirillum impatiens DSM 12546* | | **Proteobacteria GammaProteobacteria** |
| *Alkanindiges illinoisensis DSM 15370* | | **Proteobacteria GammaProteobacteria** |
| *Photobacterium profundum SS9* | | **Proteobacteria GammaProteobacteria** |
| *Frateuria terrea CGMCC 1.7053* | | **Proteobacteria GammaProteobacteria** |
| *Mariprofundus ferrooxydans M34* | | **Proteobacteria ZetaProteobacteria** |
| *RIFCSPLOWO2 2 FULL 45 22* | | **PVC Chlamydiae** |
| *Chlamydophila abortus S263* | | **PVC Chlamydiae** |
| *Lentisphaerae bacterium GWF2_52_8* | | **PVC Lentisphaerae** |
| *Candidatus Omnitrophica bacterium CG1_02_46_14* | | **PVC Omnitrophica** |
| *Planctomycetes bacterium GWB2_41_19* | | **PVC Planctomycetes** |
| *Phycisphaerae bacterium SM1_79* | | **PVC Plantomycetes** |
| *Arcanobacterium haemolyticum DSM 20595* | | **Terrabcteria Actinobacteria** |
| *Corynebacterium argentoratense DSM 44202* | | **Terrabcteria Actinobacteria** |
| *Gulosibacter molinativorax DSM 13485* | | **Terrabcteria Actinobacteria** |
| *Aestuariimicrobium kwangyangense DSM 21549* | | **Terrabcteria Actinobacteria** |
| *Bifidobacterium animalis animalis ATCC 25527* | | **Terrabcteria Actinobacteria** |
| *Eggerthella sp. YY7918* | | **Terrabcteria Actinobacteria** |
| *Rhodoluna lacicola MWH Ta8* | | **Terrabcteria Actinobacteria** |
| *Enorma massiliensis phI* | | **Terrabcteria Actinobacteria** |
| *Anaerolinea thermophila UNI-1* | | **Terrabcteria Chloroflexi** |

| | | |
|---|---|---|
| *Dehalococcoidia bacterium DG_22* | | **Terrabcteria Chloroflexi** |
| *Chloroflexi bacterium RBG_13_50_10* | | **Terrabcteria Chloroflexi** |
| *Aphanizomenon flos aquae NIES 81* | | **Terrabcteria Cyanobacteria** |
| *Crinalium epipsammum PCC 9333* | | **Terrabcteria Cyanobacteria** |
| *Oscillatoria sp. PCC 7112* | | **Terrabcteria Cyanobacteria** |
| *Stanieria cyanosphaera PCC 7437* | | **Terrabcteria Cyanobacteria** |
| *Bacillus anthracis 52 G* | | **Terrabcteria Firmicutes** |
| *Marinococcus halotolerans DSM 16375* | | **Terrabcteria Firmicutes** |
| *Fructobacillus fructosus KCTC 3544* | | **Terrabcteria Firmicutes** |
| *Streptococcus mutans GS 5* | | **Terrabcteria Firmicutes** |
| *Youngiibacter fragile 232.1* | | **Terrabcteria Firmicutes** |
| *Filifactor alocis ATCC 35896* | | **Terrabcteria Firmicutes** |
| *Orenia marismortui DSM 5156* | | **Terrabcteria Firmicutes** |
| *Veillonella parvula DSM 2008* | | **Terrabcteria Firmicutes** |
| *Melainabacteria bacterium MEL_A1* | | **Terrabcteria Melainabacteria** |
| *Candidatus Melainabacteria bacterium RIFCSPLOWO2_12_FULL_35_11* | | **Terrabcteria Melainabacteria** |
| *Mesoplasma florum W37* | | **Terrabcteria Tenericutes** |
| *Thermosynechococcus elongatus BP1* | | **Terrabcteria Cyanobacteria** |
| *Deinococcus geothermalis DSM 11300* | | **Terrabcteria Deinococcus-Thermus** |

| | | |
|---|---|---|
| *Acidobacteria bacterium RBG_16_68_9* | | **Acidobacteria** |
| *Acidobacteria bacterium RIFCSPLOWO2_02_FULL_68_18* | | **Acidobacteria** |
| *Acidobacterium sp. MP5ACTX8* | | **Acidobacteria** |
| *Holophaga foetida TMBS4 DSM 6591* | | **Acidobacteria** |
| *Candidatus Aminicenantes bacterium RBG_16_66_30* | | **Aminicenantes** |
| *Hydrogenobacter thermophilus TK 6* | | **Aquificae** |
| *Thermovibrio ammonificans HB 1* | | **Aquificae** |
| *Armatimonadetes bacterium 13_1_40CM_64_14* | | **Armatimonadetes** |
| *Bacteroides fragilis NCTC 9343* | | **Bacteroidetes** |
| *Marinilabilia salmonicolor JCM 21150* | | **Bacteroidetes** |
| *Cyclobacterium marinum DSM 745* | | **Bacteroidetes** |
| *Indibacter alkaliphilus LW1* | | **Bacteroidetes** |
| *Epilithonimonas tenax DSM 16811* | | **Bacteroidetes** |
| *Galbibacter sp. ck I2 15* | | **Bacteroidetes** |
| *Gracilimonas tropica DSM 19535* | | **Bacteroidetes** |
| *Arcticibacter svalbardensis MN12 7* | | **Bacteroidetes** |
| *Chrysiogenes arsenatis DSM 11915* | | **Chrysiogenetes** |
| *Denitrovibrio acetiphilus DSM 12809* | | **Deferribacteres** |
| *Fusobacterium nucleatum nucleatum ATCC 25586* | | **Fusobacteria** |
| *Candidatus Gracilibacteria bacterium CG1_02_38_174* | | **Gracilibacteria** |
| *Candidatus Vecturithrix granuli* | | **Modulibacteria** |
| *Nitrospinae bacterium RIFCSPLOWO2_12_FULL_47_7* | | **Nitrospinae** |
| *Nitrospirae bacterium GWC2_57_9* | | **Nitrospirae** |
| *Nitrospira defluvii* | | **Nitrospirae** |
| *Brachyspira murdochii DSM 12563* | | **Spirochaetes** |
| *Leptospira biflexa serovar Patoc strain 'Patoc 1 (Paris)'* | | **Spirochaetes** |
| *Thermanaerovibrio acidaminovorans DSM 6589* | | **Synergistetes** |

| | | |
|---|---|---|
| *Thermodesulfatator indicus CIR29812 DSM 15286* | | **Thermodesulfobacteria** |
| *Kosmotoga olearia TBF 19.5.1* | | **Thermotogae** |
| *Candidatus Wirthbacteria bacterium CG2 30_54_11* | | **Wirthbacteria** |
| *Heimdallarchaeota LC2* | | **Asgard group Candidatus Heimdallarchaeota** |
| *Heimdallarchaeota LC3* | | **Asgard group Candidatus Heimdallarchaeota** |
| *Lokiarchaeum mirabilis* | | **Asgard group Candidatus Lokiarchaeota** |
| *Odinarchaeota LCB4* | | **Asgard group Candidatus Odinarchaeota** |
| *Thorarchaeota AB25* | | **Asgard group Candidatus Thorarchaeota** |
| *Candidatus Aenigmarchaeota CG_4_10_14_3_um_filter_37_21* | **Archaea** | **DPANN group Candidatus Aenigmarchaeota** |
| *Candidatus Aenigmarchaeota archaeon CG01_land_8_20_14_3_00_37_9* | | **DPANN group Candidatus Aenigmarchaeota** |
| *Candidatus Diapherotrites archaeon CG11_big_fil_rev_8_21_14_0_20_37_9* | | **DPANN group Candidatus Diapherotrite** |
| *archaeon GW2011_AR10 (Diapherotrites archaeon AR10?)* | | **DPANN group Candidatus Diapherotrite** |
| *Candidatus Huberarchaea crystalense* | | **DPANN group Candidatus Huberarchaea** |
| *Candidatus Micrarchaeum acidiphilum ARMAN-2* | | **DPANN group Candidatus Micrarchaeota** |

| | | |
|---|---|---|
| *Candidatus Micrarchaeota archaeon A_DKE* | | **DPANN group Candidatus Micrarchaeota** |
| *Candidatus Micrarchaeota archaeon Mia14* | | **DPANN group Candidatus Micrarchaeota** |
| *Candidatus Nanosalina* | | **DPANN group Candidatus Nanohaloarchaeota** |
| *Candidatus Nanosalinarum (sp. J07AB56)* | | **DPANN group Candidatus Nanohaloarchaeota** |
| *Candidatus Nanopetramus SG9* | | **DPANN group Candidatus Nanohaloarchaeota** |
| *archaeon GW2011_AR13* | | **DPANN group Candidatus Pacearchaeota** |
| *archaeon GW2011_AR1* | | **DPANN group Candidatus Pacearchaeota** |
| *archaeon GW2011_AR6* | | **DPANN group Candidatus Pacearchaeota** |
| *Candidatus Parvarchaeum acidiphilum ARMAN-4* | | **DPANN group Candidatus Parvarchaeota** |
| *Candidatus Parvarchaeum acidophilus ARMAN-5* | | **DPANN group Candidatus Parvarchaeota** |
| *archaeon GW2011_AR15* | | **DPANN group Candidatus Woesearchaeota** |
| *archaeon GW2011_AR20* | | **DPANN group Candidatus Woesearchaeota** |
| *archaeon GW2011_AR4* | | **DPANN group Candidatus Woesearchaeota** |

| | | |
|---|---|---|
| *Nanoarchaeum equitans* | | **DPANN group Nanoarchaeota** |
| *Candidatus Nanobsidianus stetteri* | | **DPANN group Nanoarchaeota** |
| *Candidatus Nanopusillus acidilobi* | | **DPANN group Nanoarchaeota** |
| *Candidatus Altiarchaeum CG_4_9_14_0_8_um_filter_32_206* | | **DPANN group Candidatus Altiarchaeales** |
| *CG_SM1 (Alt 1) - Candidatus Altiarchaeum hamiconexum* | | **DPANN group Candidatus Altiarchaeales** |
| *WOR_SM1_SCG (Alt 2)* | | **DPANN group Candidatus Altiarchaeales** |
| *MSI_SMI (Alt 1) - Candidatus Altiarchaeum hamiconexum* | | **DPANN group Candidatus Altiarchaeales** |
| *Candidatus Altiarchaeales ex4484_2* | | **DPANN group Candidatus Altiarchaeales** |
| *Aciduliprofundum boonei (T469)* | | **Euryarchaeota Aciduliprofundum boonei** |
| *Archaeoglobus fulgidus (DSM 4304)* | | **Euryarchaeota Archaeoglobi** |
| *Archaeoglobus sulfaticallidus* | | **Euryarchaeota Archaeoglobi** |
| *Hadesarchaea archaeon YNP_N21* | | **Euryarchaeota Hadesarchaea** |
| *Hadesarchaea YNP_45* | | **Euryarchaeota Hadesarchaea** |
| *Natronomonas pharaonis (DSM 2160)* | | **Euryarchaeota Halobacteria** |
| *Haloarcula marismortui* | | **Euryarchaeota Halobacteria** |
| *Halobacterium salinarum NRC-1* | | **Euryarchaeota Halobacteria** |

| | | |
|---|---|---|
| *Halomicrobium mukohataei* | | **Euryarchaeota Halobacteria** |
| *Haloferax volcanii* | | **Euryarchaeota Halobacteria** |
| *Natronococcus amylolyticus* | | **Euryarchaeota Halobacteria** |
| *Natronobacterium gregoryi (SP2)* | | **Euryarchaeota Halobacteria** |
| *Methanobacterium bryantii* | | **Euryarchaeota Methanobacteria** |
| *Methanobacterium formicicum* | | **Euryarchaeota Methanobacteria** |
| *Methanobacterium sp. 42_16* | | **Euryarchaeota Methanobacteria** |
| *Methanobacterium subterraneum* | | **Euryarchaeota Methanobacteria** |
| *Methanothermobacter marburgensis* | | **Euryarchaeota Methanobacteria** |
| *Methanothermobacter thermautotrophicus* | | **Euryarchaeota Methanobacteria** |
| *Methanobrevibacter arboriphilus* | | **Euryarchaeota Methanobacteria** |
| *Methanobrevibacter cuticularis* | | **Euryarchaeota Methanobacteria** |
| *Methanobrevibacter filiformis* | | **Euryarchaeota Methanobacteria** |
| *Methanobrevibacter oralis JMR01* | | **Euryarchaeota Methanobacteria** |
| *Methanobrevibacter millerae* | | **Euryarchaeota Methanobacteria** |
| *Methanobrevibacter ruminantium* | | **Euryarchaeota Methanobacteria** |
| *Methanobrevibacter smithii* | | **Euryarchaeota Methanobacteria** |
| *Methanosphaera cuniculi* | | **Euryarchaeota Methanobacteria** |
| *Methanosphaera stadtmanae* | | **Euryarchaeota Methanobacteria** |

| | | |
|---|---|---|
| *Methanosphaera sp. WGK6* | | **Euryarchaeota Methanobacteria** |
| *Methanothermus fervidus* | | **Euryarchaeota Methanobacteria** |
| *Methanocaldococcus jannaschii* | | **Euryarchaeota Methanococci** |
| *Methanotorris formicicus* | | **Euryarchaeota Methanococci** |
| *Methanococcus maripaludis* | | **Euryarchaeota Methanococci** |
| *Methanothermococcus thermolithotrophicus* | | **Euryarchaeota Methanococci** |
| *Methanocella arvoryzae (Rice Cluster I)* | | **Euryarchaeota Methanomicrobia** |
| *Methanocella paludicola (SANAE)* | | **Euryarchaeota Methanomicrobia** |
| *Methanocorpusculum parvum* | | **Euryarchaeota Methanomicrobia** |
| *Methanocorpusculum labreanum* | | **Euryarchaeota Methanomicrobia** |
| *Methanoculleus bourgensis* | | **Euryarchaeota Methanomicrobia** |
| *Methanospirillum hungatei* | | **Euryarchaeota Methanomicrobia** |
| *Candidatus Methanoperedens nitroreducens (ANME-2d)* | | **Euryarchaeota Methanomicrobia** |
| *Methanothrix soehngenii* | | **Euryarchaeota Methanomicrobia** |
| *Methanothrix thermoacetophila (PT)* | | **Euryarchaeota Methanomicrobia** |
| *Methanococcoides methylutens* | | **Euryarchaeota Methanomicrobia** |
| *Methanohalophilus mahii* | | **Euryarchaeota Methanomicrobia** |
| *Methanosarcina acetivorans* | | **Euryarchaeota Methanomicrobia** |
| *Methanosarcina barkeri str. Fusaro* | | **Euryarchaeota Methanomicrobia** |

| | | |
|---|---|---|
| *Methanosarcina barkeri CM1* | | **Euryarchaeota Methanomicrobia** |
| *Methanosarcina mazei* | | **Euryarchaeota Methanomicrobia** |
| *Methanosarcina spelaei* | | **Euryarchaeota Methanomicrobia** |
| *Methermicoccus shengliensis* | | **Euryarchaeota Methanomicrobia** |
| *Methanosarcinales archaeon ex4484_138 (GoM-Arcl)* | | **Euryarchaeota Methanomicrobia** |
| *Methanosarcinales archaeon ex4572_44 (GoM-Arcl)* | | **Euryarchaeota Methanomicrobia** |
| *Candidatus Syntrophoarchaeum caldarius (ANME-2 - GoM-Arch87-2)* | | **Euryarchaeota Methanomicrobia** |
| *ANME-2 cluster archaeon HR1* | | **Euryarchaeota Methanomicrobia** |
| *ANME-1 cluster archaeon ex4572_4* | | **Euryarchaeota Methanomicrobia** |
| *Arc I group archaeon U1lsi0528_Bin055* | | **Euryarchaeota Methanomicrobia** |
| *Methanonatronarchaeum thermophilum* | | **Euryarchaeota Methanonatronarchaeia** |
| *Methanopyrus kandleri AV19* | | **Euryarchaeota Methanopyri** |
| *Methanopyrus KOL6* | | **Euryarchaeota Methanopyri** |
| *Theionarchaea archaeon DG-70-1* | | **Euryarchaeota Theionarchaea** |
| *Theionarchaea archaeon DG-70* | | **Euryarchaeota Theionarchaea** |
| *Pyrococcus furiosus* | | **Euryarchaeota Thermococci** |
| *Pyrococcus abyssi* | | **Euryarchaeota Thermococci** |
| *Pyrococcus horikoshii* | | **Euryarchaeota Thermococci** |
| *Thermococcus kodakarensis* | | **Euryarchaeota Thermococci** |

| | | |
|---|---|---|
| *Methanomassiliicoccus luminyensis* | | **Euryarchaeota Thermoplasmata** |
| *Candidatus Methanomassiliicoccus intestinalis* | | **Euryarchaeota Thermoplasmata** |
| *Candidatus Methanoplasma termitum* | | **Euryarchaeota Thermoplasmata** |
| *Acidiplasma aeolicum* | | **Euryarchaeota Thermoplasmata** |
| *Ferroplasma acidarmanus fer1* | | **Euryarchaeota Thermoplasmata** |
| *Picrophilus torridus* | | **Euryarchaeota Thermoplasmata** |
| *Thermoplasma volcanium* | | **Euryarchaeota Thermoplasmata** |
| *Thermoplasmatales archaeon SCGC AB-539-N05* | | **Euryarchaeota Thermoplasmata** |
| *Thermoplasmatales archaeon B_DKE* | | **Euryarchaeota Thermoplasmata** |
| *Cuniculiplasma sp. C_DKE* | | **Euryarchaeota Thermoplasmata** |
| *uncultured marine group II euryarchaeote* | | **Euryarchaeota unclassified** |
| *Marine group III euryarchaeote CG-Epi2* | | **Euryarchaeota unclassified** |
| *MSBL1 archaeon SCGC-AAA259E19* | | **Euryarchaeota unclassified** |
| *MCG-1* | | **TACK group Candidatus Bathyarchaeota** |
| *MCG-6* | | **TACK group Candidatus Bathyarchaeota** |
| *MCG-15* | | **TACK group Candidatus Bathyarchaeota** |
| *Candidatus Bathyarchaeota archaeon BA2* | | **TACK group Candidatus Bathyarchaeota** |

| | | |
|---|---|---|
| *Geothermarchaeota (ex4572_27)* | | **TACK group Candidatus Geothermarchaeota** |
| *Candidatus Korarchaeum cryptofilum* | | **TACK group Candidatus Korarchaeota** |
| *Methanomethylicus mesodigestum V2* | | **TACK group Candidatus Verstraetearchaeota** |
| *Methanosuratus petracarbonis V4* | | **TACK group Candidatus Verstraetearchaeota** |
| *Acidilobus saccharovorans* | | **TACK group Crenarchaeota** |
| *Aeropyrum camini* | | **TACK group Crenarchaeota** |
| *Aeropyrum pernix (K1)* | | **TACK group Crenarchaeota** |
| *Hyperthermus butylicus* | | **TACK group Crenarchaeota** |
| *Ignisphaera aggregans (DSM 17230)* | | **TACK group Crenarchaeota** |
| *Ignicoccus hospitalis* | | **TACK group Crenarchaeota** |
| *Ignicoccus islandicus (DSM 13165)* | | **TACK group Crenarchaeota** |
| *Staphylothermus marinus* | | **TACK group Crenarchaeota** |
| *Fervidicoccus fontis* | | **TACK group Crenarchaeota** |
| *Sulfolobus solfataricus* | | **TACK group Crenarchaeota** |
| *Caldivirga maquilingensis* | | **TACK group Crenarchaeota** |
| *Pyrobaculum aerophilum (str. IM2)* | | **TACK group Crenarchaeota** |
| *Thermofilum pendens* | | **TACK group Crenarchaeota** |

| | | |
|---|---|---|
| *Candidatus Caldiarchaeum subterraneum* | | **TACK group Aigarchaeota** |
| *Cenarchaeum symbiosum A* | | **TACK group Thaumarchaeota** |
| *Nitrosopumilus koreensis* | | **TACK group Thaumarchaeota** |
| *Nitrosopumilus maritimus SCM1* | | **TACK group Thaumarchaeota** |
| *Nitrosopumilus limnia* | | **TACK group Thaumarchaeota** |
| *Candidatus Nitrosomarinus catalina* | | **TACK group Thaumarchaeota** |
| *Candidatus Nitrososphaera gargensis* | | **TACK group Thaumarchaeota** |
| *Candidatus Nitrocosmicus oleophilus* | | **TACK group Thaumarchaeota** |
| *Candidatus Nitrosocaldus islandicus* | | **TACK group Thaumarchaeota** |
| *Marine group I thaumarchaeote SCGC RSA3* | | **TACK group Thaumarchaeota** |
| *Thaumarchaeota archaeon SCGC AB-539-E09* | | **TACK group Thaumarchaeota** |
| *Thaumarchaeota archaeon BS4 (Candidatus Nitrosocaldus cavascurensis)* | | **TACK group Thaumarchaeota** |
| *Thaumarchaeota Fn1* | | **TACK group Thaumarchaeota** |
| *Candidatus Marsarchaeota G2 archaeon BE_D* | | **unclassified** |
| *Candidatus Marsarchaeota G1 archaeon OSP_C* | | **unclassified** |

**Table S2: List of proteins on the hydrogenosome surface**

| Protein Accession Number | gene_ID | P_keilini protein header | function |
|---|---|---|---|
| XP_001323739.1 | TVAG_005910 | Pkeilini_DN667_c0_g2_i1.p1 | 50S-ribosomal-protein-L2,-putative |
| XP_001323765.1 | TVAG_006170 | Pkeilini_DN989_c0_g1_i1.p1 | 60S-ribosomal-protein-L19,-putative |
| XP_001323773.1 | TVAG_006250 | Pkeilini_DN915_c0_g1_i1.p1 | 30S-ribosomal-protein-S8,-putative |
| XP_001311950.1 | TVAG_008680 | Pkeilini_DN1020_c0_g1_i1.p1 | tubulin-epsilon-chain,-putative |
| XP_001582527.1 | TVAG_013060 | Pkeilini_DN4044_c0_g1_i1.p1 | 60S-ribosomal-protein-L3,-putative |
| XP_001582635.1 | TVAG_014160 | Pkeilini_DN645_c0_g1_i1.p2 | 60S-ribosomal-protein-L12,-putative |
| XP_001329886.1 | TVAG_015800 | Pkeilini_DN460_c0_g1_i2.p1 | 60S-ribosomal-protein-L23,-putative |
| XP_001317481.1 | TVAG_020040 | Pkeilini_DN3914_c0_g3_i1.p1 | ribosomal-protein-S9,-putative |
| XP_001314734.1 | TVAG_020480 | Pkeilini_DN508_c0_g2_i1.p1 | 40S-ribosomal-protein-S18,-putative |
| XP_001300897.1 | TVAG_033590 | Pkeilini_DN670_c0_g1_i1.p1 | 40S-ribosomal-protein-S6,-putative |
| XP_001326922.1 | TVAG_038050 | Pkeilini_DN414_c0_g1_i2.p1 | 50S-ribosomal-protein-L24p,-putative |
| XP_001315155.1 | TVAG_040820 | Pkeilini_DN16934_c0_g1_i1.p2 | 40S-ribosomal-protein-S17,-putative |
| XP_001301101.1 | TVAG_041350 | Pkeilini_DN3310_c1_g1_i1.p1 | 30S-40S-ribosomal-protein,-putative |
| XP_001312328.1 | TVAG_043060 | Pkeilini_DN163620_c0_g1_i1.p1 | fructose-bisphosphate-aldolase,-putative |
| XP_001315627.1 | TVAG_043500 | Pkeilini_DN242_c0_g2_i1.p1 | enolase,-putative |
| XP_001314691.1 | TVAG_044510 | Pkeilini_DN842_c0_g2_i4.p1 | heat-shock-protein-70-(HSP70)-4,-putative |
| XP_001330678.1 | TVAG_044560 | Pkeilini_DN374_c0_g1_i1.p1 | 50S-ribosomal-protein-L5p,-putative |
| XP_001323300.1 | TVAG_045010 | Pkeilini_DN206053_c0_g1_i1.p1 | glucokinase,-putative |
| XP_001308576.1 | TVAG_045340 | Pkeilini_DN420_c0_g1_i1.p1 | polyadenylate-binding-protein,-putative |

| | | | |
|---|---|---|---|
| XP_001298952.1 | TVAG_047460 | Pkeilini_DN207414_c0_g1_i1.p1 | 40S-ribosomal-protein-S3a,-putative |
| XP_001321104.1 | TVAG_051160 | Pkeilini_DN182683_c0_g1_i1.p1 | 60S-acidic-ribosomal-protein-P0,-putative |
| XP_001306738.1 | TVAG_054130 | Pkeilini_DN92336_c3_g1_i1.p1 | 60S-ribosomal-protein-L7,-putative |
| XP_001330619.1 | TVAG_054500 | Pkeilini_DN198864_c0_g1_i1.p1 | 50S-ribosomal-protein-L6p,-putative |
| XP_001308434.1 | TVAG_054830 | Pkeilini_DN184234_c0_g1_i1.p1 | phosphoglucomutase,-putative |
| XP_001323508.1 | TVAG_061890 | Pkeilini_DN161037_c0_g1_i1.p1 | 60S-ribosomal-protein-L18a,-putative |
| XP_001323512.1 | TVAG_061930 | Pkeilini_DN218798_c0_g1_i1.p1 | glucose-6-phosphate-isomerase,-putative |
| XP_001320140.1 | TVAG_064640 | Pkeilini_DN170420_c1_g1_i4.p1 | ribosomal-protein-L5,-putative |
| XP_001318569.1 | TVAG_066030 | Pkeilini_DN1889_c0_g1_i1.p1 | 40S-ribosomal-protein-S8,-putative |
| XP_001328958.1 | TVAG_067400 | Pkeilini_DN828_c0_g1_i2.p1 | elongation-factor-1-alpha,-putative |
| XP_001584514.1 | TVAG_071920 | Pkeilini_DN103_c0_g1_i1.p1 | 40S-ribosomal-protein-S23,-putative |
| XP_001321243.1 | TVAG_073860 | Pkeilini_DN2499_c0_g3_i2.p1 | phosphoenolpyruvate-protein-phosphotransferase,-putative |
| XP_001324843.1 | TVAG_074480 | Pkeilini_DN283_c0_g1_i2.p1 | 60S-ribosomal-protein-L10a,-putative |
| XP_001324856.1 | TVAG_074610 | Pkeilini_DN1754_c0_g1_i2.p1 | 60S-ribosomal-protein-L10,-putative |
| XP_001320891.1 | TVAG_079260 | Pkeilini_DN9_c0_g1_i2.p1 | phosphofructokinase,-putative |
| XP_001579459.1 | TVAG_083260 | Pkeilini_DN570_c0_g4_i1.p1 | 60S-ribosomal-protein-L17,-putative |
| XP_001318159.1 | TVAG_087140 | Pkeilini_DN173700_c0_g1_i1.p1 | arp2/3,-putative |
| XP_001304619.1 | TVAG_092750 | Pkeilini_DN690_c0_g1_i1.p1 | glucokinase |
| XP_001303733.1 | TVAG_094720 | Pkeilini_DN605_c0_g1_i1.p2 | 60S-ribosomal-protein-L34,-putative |
| XP_001321933.1 | TVAG_098450 | Pkeilini_DN218705_c0_g1_i1.p1 | 40S-ribosomal-protein-S4,-putative |

| | | | |
|---|---|---|---|
| XP_001303193.1 | TVAG_099490 | Pkeilini_DN184416_c0_g1_i1.p1 | glucose-kinase,-putative |
| XP_001580437.1 | TVAG_101690 | Pkeilini_DN165121_c0_g1_i1.p1 | 60S-ribosomal-protein-L24,-putative |
| XP_001324008.1 | TVAG_106800 | Pkeilini_DN1152_c0_g1_i1.p1 | 30S-ribosomal-protein-S3,-putative |
| XP_001581388.1 | TVAG_110140 | Pkeilini_DN167_c0_g3_i5.p2 | ubiquitin,-putative |
| XP_001300910.1 | TVAG_112230 | Pkeilini_DN79301_c0_g2_i1.p1 | 60S-ribosomal-protein-L13,-putative |
| XP_001579013.1 | TVAG_113710 | Pkeilini_DN54853_c0_g1_i1.p2 | phosphoglycerate-mutase,-putative |
| XP_001324949.1 | TVAG_117480 | Pkeilini_DN330_c0_g1_i3.p1 | 30S-ribosomal-protein-S11,-putative |
| XP_001276845.1 | TVAG_119330 | Pkeilini_DN19_c0_g6_i1.p1 | 60S-ribosomal-protein-L21,-putative |
| XP_001276908.1 | TVAG_119970 | Pkeilini_DN208907_c0_g1_i1.p1 | conserved-hypothetical-protein |
| XP_001276929.1 | TVAG_120180 | Pkeilini_DN9217_c0_g1_i1.p1 | 40S-ribosomal-protein-S10,-putative |
| XP_001277020.1 | TVAG_121100 | Pkeilini_DN426_c0_g1_i2.p1 | 60S-ribosomal-protein-L18,-putative |
| XP_001322974.1 | TVAG_121550 | Pkeilini_DN1040_c0_g1_i1.p1 | 60S-ribosomal-protein-L27a,-putative |
| XP_001324688.1 | TVAG_128790 | Pkeilini_DN68345_c0_g1_i2.p1 | 60S-ribosomal-protein-L4,-putative |
| XP_001324775.1 | TVAG_139320 | Pkeilini_DN43315_c0_g1_i1.p1 | heat-shock-protein,-putative |
| XP_001320113.1 | TVAG_142440 | Pkeilini_DN3544_c0_g1_i2.p1 | 40S-ribosomal-protein-S2,-putative |
| XP_001579934.1 | TVAG_146910 | Pkeilini_DN183802_c0_g1_i1.p1 | glyceraldehyde-3-phosphate-dehydrogenase,-putative |
| XP_001309550.1 | TVAG_148950 | Pkeilini_DN639_c0_g1_i1.p1 | 50S-ribosomal-protein-L15e,-putative |
| XP_001309564.1 | TVAG_149090 | Pkeilini_DN976_c0_g2_i4.p1 | actin,-putative |
| XP_001303635.1 | TVAG_152720 | Pkeilini_DN330_c0_g1_i3.p1 | 40S-ribosomal-protein-S14,-putative |
| XP_001317545.1 | TVAG_153560 | Pkeilini_DN358_c0_g1_i4.p1 | heat-shock-protein,-putative |

| | | | |
|---|---|---|---|
| XP_001313895.1 | TVAG_154680 | Pkeilini_DN5492_c0_g1_i1.p1 | conserved-hypothetical-protein |
| XP_001310739.1 | TVAG_157940 | Pkeilini_DN5492_c0_g1_i1.p1 | conserved-hypothetical-protein |
| XP_001580742.1 | TVAG_178000 | Pkeilini_DN52_c0_g1_i2.p1 | 60S-ribosomal-protein-L23a,-putative |
| XP_001583987.1 | TVAG_182370 | Pkeilini_DN69_c0_g3_i2.p1 | chaperonin-containing-t-complex-protein-1,-gamma-subunit,-tcpg,-putative |
| XP_001584002.1 | TVAG_182520 | Pkeilini_DN79301_c0_g2_i1.p1 | 60S-ribosomal-protein-L13,-putative |
| XP_001580136.1 | TVAG_190450 | Pkeilini_DN1019_c0_g1_i1.p1 | kakapo,-putative |
| XP_001320814.1 | TVAG_191140 | Pkeilini_DN528_c0_g1_i1.p1 | conserved-hypothetical-protein |
| XP_001581272.1 | TVAG_192620 | Pkeilini_DN206419_c0_g1_i1.p1 | actin-depolymerizing-factor,-putative |
| XP_001308024.1 | TVAG_204360 | Pkeilini_DN168_c0_g2_i1.p1 | malate-dehydrogenase,-putative |
| XP_001308025.1 | TVAG_204370 | Pkeilini_DN206053_c0_g1_i1.p1 | glucokinase,-putative |
| XP_001303801.1 | TVAG_205910 | Pkeilini_DN184234_c0_g1_i1.p1 | phosphoglucomutase,-putative |
| XP_001319786.1 | TVAG_212020 | Pkeilini_DN5548_c1_g1_i1.p1 | transketolase,-putative |
| XP_001325073.1 | TVAG_222040 | Pkeilini_DN5492_c0_g1_i1.p1 | conserved-hypothetical-protein |
| XP_001310373.1 | TVAG_226870 | Pkeilini_DN5492_c0_g1_i1.p1 | conserved-hypothetical-protein |
| XP_001303253.1 | TVAG_234160 | Pkeilini_DN219244_c0_g1_i1.p1 | arp2/3-complex-20-kD-subunit,-putative |
| XP_001581543.1 | TVAG_239310 | Pkeilini_DN1019_c0_g2_i2.p1 | bollus-pemphigoid-antigen,-putative |
| XP_001320867.1 | TVAG_240050 | Pkeilini_DN516_c0_g1_i1.p1 | 40S-ribosomal-protein-sa,-putative |
| XP_001323701.1 | TVAG_248450 | Pkeilini_DN1700_c0_g1_i1.p1 | peptidyl-tRNA-hydrolase,-putative |
| XP_001579239.1 | TVAG_253650 | Pkeilini_DN168_c0_g2_i1.p1 | malate-dehydrogenase,-putative |
| XP_001323043.1 | TVAG_258220 | Pkeilini_DN7684_c0_g1_i1.p1 | glycosyltransferase,-putative |

| | | | |
|---|---|---|---|
| XP_001317828.1 | TVAG_263740 | Pkeilini_DN242_c0_g1_i2.p1 | enolase,-putative |
| XP_001313821.1 | TVAG_265950 | Pkeilini_DN56722_c0_g1_i1.p1 | 60S-ribosomal-protein-L32,-putative |
| XP_001579758.1 | TVAG_268050 | Pkeilini_DN220_c0_g1_i1.p1 | phosphoglycerate-kinase,-putative |
| XP_001330352.1 | TVAG_272970 | Pkeilini_DN362_c0_g1_i1.p1 | 40S-ribosomal-protein-S24,-putative |
| XP_001321781.1 | TVAG_276310 | Pkeilini_DN217279_c1_g1_i1.p1 | starch-branching-enzyme-II,-putative |
| XP_001321791.1 | TVAG_276410 | Pkeilini_DN3469_c0_g2_i1.p1 | translation-elongation-factor,-putative |
| XP_001304599.1 | TVAG_277390 | Pkeilini_DN160592_c0_g1_i1.p2 | peptidyl-prolyl-cis-trans-isomerase-A,-ppia,-putative |
| XP_001582074.1 | TVAG_282580 | Pkeilini_DN52882_c0_g1_i1.p1 | conserved-hypothetical-protein |
| XP_001582118.1 | TVAG_283020 | Pkeilini_DN587_c0_g1_i1.p1 | initiation-factor-5A,-putative |
| XP_001330357.1 | TVAG_292580 | Pkeilini_DN16084_c3_g1_i1.p1 | immunophilin,-putative |
| XP_001307577.1 | TVAG_293770 | Pkeilini_DN160642_c0_g1_i1.p1 | phosphofructokinase,-putative |
| XP_001315498.1 | TVAG_299380 | Pkeilini_DN330_c0_g1_i3.p1 | 30S-ribosomal-protein-S11,-putative |
| XP_001311055.1 | TVAG_300000 | Pkeilini_DN163620_c0_g1_i1.p1 | fructose-bisphosphate-aldolase,-putative |
| XP_001304306.1 | TVAG_319220 | Pkeilini_DN362_c0_g2_i1.p1 | 40S-ribosomal-protein-S24,-putative |
| XP_001322282.1 | TVAG_329460 | Pkeilini_DN242_c0_g2_i1.p1 | enolase,-putative |
| XP_001317525.1 | TVAG_336940 | Pkeilini_DN690_c0_g1_i1.p1 | glucokinase,-putative |
| XP_001319342.1 | TVAG_342830 | Pkeilini_DN330_c0_g1_i3.p1 | 40S-ribosomal-protein-S14/30S-ribosomal-protein-S11,-putative |
| XP_001328722.1 | TVAG_348090 | Pkeilini_DN207414_c0_g1_i1.p1 | 40S-ribosomal-protein-S3a,-putative |
| XP_001328746.1 | TVAG_348330 | Pkeilini_DN921_c0_g2_i1.p1 | glycogen-phosphorylase,-putative |
| XP_001326306.1 | TVAG_351310 | Pkeilini_DN351_c0_g1_i1.p1 | plastin,-putative |

| | | | |
|---|---|---|---|
| XP_001308720.1 | TVAG_354020 | Pkeilini_DN160502_c0_g1_i1.p1 | centractin,-putative |
| XP_001307153.1 | TVAG_360700 | Pkeilini_DN163620_c0_g1_i1.p1 | fructose-bisphosphate-aldolase,-putative |
| XP_001307814.1 | TVAG_370550 | Pkeilini_DN670_c0_g1_i1.p1 | 40S-ribosomal-protein-S6,-putative |
| XP_001305702.1 | TVAG_373720 | Pkeilini_DN2724_c0_g1_i10.p2 | pyruvate-kinase,-putative |
| XP_001306345.1 | TVAG_376130 | Pkeilini_DN16874_c0_g1_i1.p3 | gelsolin,-putative |
| XP_001304373.1 | TVAG_380910 | Pkeilini_DN600_c0_g3_i1.p1 | DEAD-box-ATP-dependent-RNA-helicase,-putative |
| XP_001295243.1 | TVAG_381030 | Pkeilini_DN382_c0_g1_i1.p1 | conserved-hypothetical-protein |
| XP_001302740.1 | TVAG_381690 | Pkeilini_DN208349_c0_g1_i1.p1 | NAD-dependent-epimerase/dehydratase,-putative |
| XP_001314248.1 | TVAG_383940 | Pkeilini_DN220_c0_g1_i1.p1 | phosphoglycerate-kinase,-putative |
| XP_001581778.1 | TVAG_391760 | Pkeilini_DN450_c0_g1_i1.p2 | phosphofructokinase,-putative |
| XP_001327217.1 | TVAG_397250 | Pkeilini_DN690_c0_g1_i1.p1 | glucokinase,-putative |
| XP_001328502.1 | TVAG_423320 | Pkeilini_DN79301_c0_g2_i1.p1 | 60S-ribosomal-protein-L13,-putative |
| XP_001325139.1 | TVAG_430830 | Pkeilini_DN9_c0_g1_i2.p1 | phosphofructokinase,-putative |
| XP_001302863.1 | TVAG_435000 | Pkeilini_DN16084_c3_g1_i1.p1 | conserved-hypothetical-protein |
| XP_001313948.1 | TVAG_442070 | Pkeilini_DN184416_c0_g1_i1.p1 | glucose-kinase,-putative |
| XP_001579592.1 | TVAG_462920 | Pkeilini_DN160642_c0_g1_i1.p1 | phosphofructokinase,-putative |
| XP_001325501.1 | TVAG_464120 | Pkeilini_DN330_c0_g1_i1.p1 | 30S-ribosomal-protein-S11,-putative |
| XP_001325506.1 | TVAG_464170 | Pkeilini_DN242_c0_g2_i1.p1 | enolase,-putative |
| XP_001322192.1 | TVAG_482430 | Pkeilini_DN426_c0_g1_i2.p1 | 60S-ribosomal-protein-L18,-putative |
| XP_001322532.1 | TVAG_491670 | Pkeilini_DN207881_c0_g1_i1.p1 | malic-enzyme,-putative |

| | | | |
|---|---|---|---|
| XP_001305512.1 | TVAG_496160 | Pkeilini_DN211320_c0_g1_i1.p1 | phosphofructokinase,-putative |

**Table S3: List of putative membrane proteins, enzymes and hypothetical proteins identified in the hydrogenosome**

| Protein Accession Number | gene_ID | P_keilini protein header | function |
|---|---|---|---|
| XP_001584373.1 | TVAG_070500 | Pkeilini_DN58243_c4_g1_i1.p1 | Rab7g-protein,-putative |
| XP_001584268.1 | TVAG_185900 | Pkeilini_DN211195_c0_g1_i1.p1 | conserved-hypothetical-protein |
| XP_001584214.1 | TVAG_185340 | Pkeilini_DN171252_c0_g1_i1.p1 | Rab32,-putative |
| XP_001584210.1 | TVAG_185300 | Pkeilini_DN25505_c0_g1_i1.p1 | Rab11,-putative |
| XP_001584188.1 | TVAG_185080 | Pkeilini_DN175067_c0_g1_i1.p1 | ran,-putative |
| XP_001584134.1 | TVAG_183850 | Pkeilini_DN5301_c0_g1_i1.p1 | conserved-hypothetical-protein |
| XP_001584128.1 | TVAG_183790 | Pkeilini_DN568_c0_g1_i1.p1 | malic-enzyme,-putative |
| XP_001584099.1 | TVAG_183500 | Pkeilini_DN541_c0_g1_i1.p1 | succinate-thiokinase-¿-subunit |
| XP_001584080.1 | TVAG_183300 | Pkeilini_DN194471_c0_g1_i1.p1 | aminotransferase-class-V,-putative |
| XP_001584012.1 | TVAG_182620 | Pkeilini_DN13155_c3_g1_i1.p1 | nitrate,-fromate,-iron-dehydrogenase,-putative |
| XP_001583906.1 | TVAG_076670 | Pkeilini_DN1671_c1_g1_i1.p1 | small-GTPase-RAB,-putative |
| XP_001583890.1 | TVAG_076510 | Pkeilini_DN175_c0_g1_i1.p1 | 2-amino-3-ketobutyrate-coenzyme-A-ligase,-putative |
| XP_001583862.1 | TVAG_076230 | Pkeilini_DN584_c0_g1_i1.p1 | nucleotide-binding-protein,-putative |
| XP_001583785.1 | TVAG_075460 | Pkeilini_DN206_c3_g1_i1.p1 | Rab5b,-putative |

| | | | |
|---|---|---|---|
| XP_001583584.1 | TVAG_036230 | Pkeilini_DN26_c4_g1_i1.p1 | RAB,-putative |
| XP_001583562.1 | TVAG_036010 | Pkeilini_DN160973_c0_g1_i1.p1 | A-type-flavoprotein |
| XP_001583303.1 | TVAG_377960 | Pkeilini_DN26_c4_g1_i1.p1 | Rab8,-putative |
| XP_001583118.1 | TVAG_093060 | Pkeilini_DN877_c0_g2_i1.p1 | Rabx30-protein,-putative |
| XP_001583042.1 | TVAG_456910 | Pkeilini_DN941_c2_g5_i1.p1 | Rabx32-protein,-putative |
| XP_001583028.1 | TVAG_456770 | Pkeilini_DN419_c0_g1_i1.p2 | iron-sulfur-cluster-assembly-protein,-putative |
| XP_001582848.1 | TVAG_249220 | Pkeilini_DN953_c0_g3_i1.p1 | Rab2,-putative |
| XP_001582728.1 | TVAG_237680 | Pkeilini_DN208767_c1_g1_i1.p1 | ADP,ATP-carrier-protein,-putative |
| XP_001582674.1 | TVAG_237140 | Pkeilini_DN842_c0_g4_i1.p1 | heat-shock-protein,-putative |
| XP_001582391.1 | TVAG_198430 | Pkeilini_DN195283_c0_g1_i1.p1 | 14-3-3-protein-sigma,-gamma,-zeta,-beta/alpha,-putative |
| XP_001582383.1 | TVAG_198350 | Pkeilini_DN6701_c0_g1_i1.p2 | hypothetical-protein |
| XP_001582360.1 | TVAG_198110 | Pkeilini_DN688_c0_g3_i1.p1 | pyruvate-flavodoxin-oxidoreductase,-putative |
| XP_001582336.1 | TVAG_167250 | Pkeilini_DN392_c0_g1_i1.p1 | chaperonin,-putative |
| XP_001582335.1 | TVAG_167240 | Pkeilini_DN16677_c0_g1_i1.p1 | conserved-hypothetical-protein |
| XP_001582155.1 | TVAG_283380 | Pkeilini_DN2717_c0_g1_i1.p1 | conserved-hypothetical-protein |
| XP_001581795.1 | TVAG_391930 | Pkeilini_DN184428_c0_g1_i1.p1 | RAB,-putative |
| XP_001581497.1 | TVAG_238830 | Pkeilini_DN568_c0_g1_i1.p1 | malic-enzyme,-putative |
| XP_001581254.1 | TVAG_192440 | Pkeilini_DN941_c3_g1_i1.p1 | RAC,-putative |
| XP_001580948.1 | TVAG_402160 | Pkeilini_DN21606_c0_g1_i1.p1 | guanine-nucleotide-exchange-factor,-putative |
| XP_001580911.1 | TVAG_130330 | Pkeilini_DN22962_c0_g1_i1.p1 | conserved-hypothetical-protein |

| | | | |
|---|---|---|---|
| XP_001580780.1 | TVAG_178380 | Pkeilini_DN160711_c0_g1_i1.p1 | conserved-hypothetical-protein |
| XP_001580752.1 | TVAG_178100 | Pkeilini_DN194385_c0_g1_i1.p1 | hypothetical-protein |
| XP_001580745.1 | TVAG_178030 | Pkeilini_DN160251_c0_g1_i1.p1 | plasma-membrane-calcium-transporting-ATPase,-putative |
| XP_001580733.1 | TVAG_177910 | Pkeilini_DN1508_c0_g1_i1.p1 | hypothetical-protein |
| XP_001580703.1 | TVAG_433130 | Pkeilini_DN285_c1_g1_i1.p1 | heat-shock-protein,-putative |
| XP_001580655.1 | TVAG_432650 | Pkeilini_DN198613_c0_g1_i1.p1 | nitrogen-fixation-protein-nifu,-putative |
| XP_001580601.1 | TVAG_228780 | Pkeilini_DN501_c0_g1_i1.p1 | alcohol-dehydrogenase,-putative |
| XP_001580529.1 | TVAG_136740 | Pkeilini_DN184428_c0_g1_i1.p1 | RAB,-putative |
| XP_001580481.1 | TVAG_136260 | Pkeilini_DN26_c5_g1_i1.p1 | GTP-binding-protein-ypt10,-putative |
| XP_001580416.1 | TVAG_101480 | Pkeilini_DN12181_c0_g1_i1.p1 | RAB,-putative |
| XP_001580394.1 | TVAG_101260 | Pkeilini_DN13976_c0_g1_i1.p1 | Rab32,-putative |
| XP_001580191.1 | TVAG_214300 | Pkeilini_DN15514_c0_g1_i1.p2 | conserved-hypothetical-protein |
| XP_001580148.1 | TVAG_190580 | Pkeilini_DN58662_c0_g1_i1.p1 | Clan-MG,-familly-M24,-aminopeptidase-P-like-metallopeptidase |
| XP_001580142.1 | TVAG_190510 | Pkeilini_DN4576_c0_g1_i1.p1 | GTP-binding-protein-rit,-putative |
| XP_001580044.1 | TVAG_247370 | Pkeilini_DN184336_c0_g1_i1.p1 | conserved-hypothetical-protein |
| XP_001579755.1 | TVAG_268020 | Pkeilini_DN194634_c1_g1_i1.p1 | aspartate-aminotransferase,-putative |
| XP_001579748.1 | TVAG_267950 | Pkeilini_DN207024_c0_g1_i1.p1 | lysyl-tRNA-synthetase,-putative |
| XP_001579740.1 | TVAG_267870 | Pkeilini_DN269_c0_g1_i2.p1 | malic-enzyme,-putative |
| XP_001579545.1 | TVAG_462450 | Pkeilini_DN676_c0_g2_i1.p1 | RAB,-putative |
| XP_001579538.1 | TVAG_462370 | Pkeilini_DN14322_c0_g1_i3.p1 | Rabx19-protein,-putative |

| | | | |
|---|---|---|---|
| XP_001579477.1 | TVAG_083440 | Pkeilini_DN112439_c0_g1_i1.p1 | conserved-hypothetical-protein |
| XP_001579475.1 | TVAG_083420 | Pkeilini_DN147688_c0_g2_i1.p1 | conserved-hypothetical-protein |
| XP_001579216.1 | TVAG_122960 | Pkeilini_DN52162_c0_g1_i1.p1 | conserved-hypothetical-protein |
| XP_001579154.1 | TVAG_122340 | Pkeilini_DN1085_c1_g2_i1.p1 | RAB,-putative |
| XP_001579147.1 | TVAG_122270 | Pkeilini_DN3020_c0_g2_i1.p1 | RAB,-putative |
| XP_001579030.1 | TVAG_113880 | Pkeilini_DN13377_c0_g1_i1.p1 | conserved-hypothetical-protein |
| XP_001579029.1 | TVAG_113870 | Pkeilini_DN182694_c0_g1_i1.p1 | Acetyl-CoA-hydrolase,-putative |
| XP_001578953.1 | TVAG_225930 | Pkeilini_DN28069_c0_g1_i1.p1 | conserved-hypothetical-protein |
| XP_001276882.1 | TVAG_119710 | Pkeilini_DN22730_c0_g1_i1.p1 | Clan-ME,-family-M16,-insulinase-like-metallopeptidase |
| XP_001330794.1 | TVAG_320200 | Pkeilini_DN3462_c0_g2_i1.p1 | Rab5b,-putative |
| XP_001330618.1 | TVAG_054490 | Pkeilini_DN201059_c0_g1_i1.p1 | Tryptophanase,-putative |
| XP_001330504.1 | TVAG_383530 | Pkeilini_DN177099_c0_g1_i1.p1 | RAB,-putative |
| XP_001330829.1 | TVAG_217400 | Pkeilini_DN211716_c0_g1_i1.p1 | conserved-hypothetical-protein |
| XP_001330450.1 | TVAG_047890 | Pkeilini_DN626_c0_g1_i1.p1 | succinate-thiokinase-a-subunit |
| XP_001314415.1 | TVAG_090060 | Pkeilini_DN941_c2_g4_i1.p2 | GTP-binding-protein-Rab2,-putative |
| XP_001330332.1 | TVAG_272760 | Pkeilini_DN218827_c0_g1_i1.p1 | glutathione-reductase,-putative |
| XP_001330320.1 | TVAG_217870 | Pkeilini_DN19306_c0_g1_i1.p2 | nucleotide-binding-protein,-putative |
| XP_001330242.1 | TVAG_325080 | Pkeilini_DN183480_c0_g1_i1.p1 | conserved-hypothetical-protein |
| XP_001330232.1 | TVAG_324980 | Pkeilini_DN8882_c0_g2_i2.p2 | ATP-synthase,-putative |
| XP_001330176.1 | TVAG_395550 | Pkeilini_DN182694_c0_g1_i1.p1 | Acetyl-CoA-hydrolase,-putative |

| | | | |
|---|---|---|---|
| XP_001329730.1 | TVAG_454230 | Pkeilini_DN184377_c0_g1_i1.p1 | Rab15,-13,-10,-1,-35,-5,-and,-putative |
| XP_001329309.1 | TVAG_297650 | Pkeilini_DN601_c0_g1_i1.p1 | grpe-protein,-putative |
| XP_001329276.1 | TVAG_297320 | Pkeilini_DN26_c4_g1_i1.p1 | small-GTPase-rabi,-putative |
| XP_001329126.1 | TVAG_150540 | Pkeilini_DN58243_c4_g1_i1.p1 | RAB-2,4,14,-putative |
| XP_001329022.1 | TVAG_447580 | Pkeilini_DN69533_c0_g1_i1.p1 | conserved-hypothetical-protein |
| XP_001328863.1 | TVAG_434770 | Pkeilini_DN1107_c0_g1_i1.p1 | Rab5b,-putative |
| XP_001328770.1 | TVAG_348580 | Pkeilini_DN1522_c0_g1_i1.p1 | Rabx26-protein,-putative |
| XP_001328523.1 | TVAG_423530 | Pkeilini_DN4238_c0_g1_i1.p2 | conserved-hypothetical-protein |
| XP_001328415.1 | TVAG_340860 | Pkeilini_DN12251_c0_g1_i1.p2 | Rab7g-protein,-putative |
| XP_001328285.1 | TVAG_278280 | Pkeilini_DN26_c1_g1_i1.p2 | Rabx22-protein,-putative |
| XP_001328194.1 | TVAG_262210 | Pkeilini_DN172140_c0_g1_i1.p1 | tricarboxylate-transport-protein,-putative |
| XP_001328129.1 | TVAG_165340 | Pkeilini_DN626_c0_g1_i1.p1 | succinate-thiokinase-a-subunit |
| XP_001328031.1 | TVAG_159810 | Pkeilini_DN953_c0_g5_i1.p2 | small-GTPase-RAB,-putative |
| XP_001328023.1 | TVAG_159730 | Pkeilini_DN206_c0_g2_i2.p1 | Rab78,-putative |
| XP_001327882.1 | TVAG_405730 | Pkeilini_DN7495_c0_g1_i1.p1 | Rabx37-protein,-putative |
| XP_001327427.1 | TVAG_201980 | Pkeilini_DN7689_c0_g1_i1.p1 | Rab15,-13,-10,-1,-35,-5,-and,-putative |
| XP_001327371.1 | TVAG_392650 | Pkeilini_DN781_c0_g1_i1.p1 | conserved-hypothetical-protein |
| XP_001327338.1 | TVAG_392320 | Pkeilini_DN199807_c0_g1_i1.p1 | groes-chaperonin,-putative |
| XP_001327242.1 | TVAG_019190 | Pkeilini_DN75143_c0_g2_i1.p1 | chaperone-protein-DNAj,-putative |
| XP_001327227.1 | TVAG_397350 | Pkeilini_DN160922_c0_g1_i1.p1 | Rab9,-putative |

| | | | |
|---|---|---|---|
| XP_001326968.1 | TVAG_038530 | Pkeilini_DN196128_c0_g1_i1.p1 | ornithine-carbamoyltransferase,-putative |
| XP_001326958.1 | TVAG_038420 | Pkeilini_DN5113_c0_g1_i1.p1 | conserved-hypothetical-protein |
| XP_001326942.1 | TVAG_038250 | Pkeilini_DN900_c0_g1_i1.p2 | Rab12,-putative |
| XP_001326936.1 | TVAG_038190 | Pkeilini_DN8719_c0_g1_i1.p1 | small-GTPase-rabi,-putative |
| XP_001326908.1 | TVAG_461020 | Pkeilini_DN3412_c0_g1_i1.p1 | ABC-transporter,-putative |
| XP_001326833.1 | TVAG_393370 | Pkeilini_DN196009_c0_g1_i1.p1 | Rab23,-putative |
| XP_001326755.1 | TVAG_388650 | Pkeilini_DN30752_c0_g1_i1.p1 | serine-palmitoyltransferase-I,-putative |
| XP_001326629.1 | TVAG_255980 | Pkeilini_DN9830_c0_g1_i1.p1 | conserved-hypothetical-protein |
| XP_001326421.1 | TVAG_373310 | Pkeilini_DN85431_c0_g1_i1.p1 | RAB,-putative |
| XP_001326329.1 | TVAG_351540 | Pkeilini_DN25815_c0_g1_i1.p1 | 2,4-dienoyl-CoA-reductase-[NADPH],-putative |
| XP_001326325.1 | TVAG_351500 | Pkeilini_DN196009_c0_g1_i1.p1 | Rab7,-putative |
| XP_001326307.1 | TVAG_351320 | Pkeilini_DN17_c0_g2_i1.p1 | purine-nucleoside-phosphorylase,-putative |
| XP_001326088.1 | TVAG_044270 | Pkeilini_DN494_c1_g1_i1.p1 | RAB,-putative |
| XP_001326029.1 | TVAG_468220 | Pkeilini_DN13197_c0_g1_i1.p2 | conserved-hypothetical-protein |
| XP_001325929.1 | TVAG_371800 | Pkeilini_DN80221_c1_g1_i1.p1 | conserved-hypothetical-protein |
| XP_001325705.1 | TVAG_206500 | Pkeilini_DN3171_c0_g1_i1.p1 | Hydroxylamine-reductase,-putative |
| XP_001325649.1 | TVAG_424580 | Pkeilini_DN175264_c0_g1_i1.p1 | mevalonate-kinase,-putative |
| XP_001325272.1 | TVAG_212310 | Pkeilini_DN12195_c0_g1_i1.p1 | Rabx21-protein,-putative |
| XP_001325171.1 | TVAG_404940 | Pkeilini_DN2224_c0_g1_i1.p1 | RAB,-putative |
| XP_001324855.1 | TVAG_074600 | Pkeilini_DN195601_c0_g1_i1.p1 | tyrosine-aminotransferase,-putative |

| | | | |
|---|---|---|---|
| XP_001324836.1 | TVAG_074410 | Pkeilini_DN1405_c0_g2_i1.p1 | Rabx41-protein,-putative |
| XP_001324773.1 | TVAG_139300 | Pkeilini_DN111_c0_g1_i1.p1 | phosphoenolpyruvate-carboxykinase,-putative |
| XP_001324770.1 | TVAG_139270 | Pkeilini_DN26_c1_g1_i1.p2 | Rab10,-putative |
| XP_001324695.1 | TVAG_128860 | Pkeilini_DN211282_c0_g1_i1.p1 | RAB-18,-putative |
| XP_001324545.1 | TVAG_161280 | Pkeilini_DN494_c0_g3_i2.p1 | Rab32,-putative |
| XP_001324510.1 | TVAG_160930 | Pkeilini_DN15873_c0_g1_i1.p1 | Periplasmic-[Fe]-hydrogenase,-putative |
| XP_001324271.1 | TVAG_038950 | Pkeilini_DN941_c0_g4_i1.p1 | Rabx3-protein,-putative |
| XP_001324262.1 | TVAG_038850 | Pkeilini_DN184336_c0_g1_i1.p1 | conserved-hypothetical-protein |
| XP_001324197.1 | TVAG_271570 | Pkeilini_DN50689_c0_g1_i1.p1 | equilibrative-nucleoside-transporter,-putative |
| XP_001324090.1 | TVAG_362470 | Pkeilini_DN7689_c0_g1_i1.p1 | Rab17,-putative |
| XP_001323999.1 | TVAG_106710 | Pkeilini_DN726_c0_g3_i1.p1 | conserved-hypothetical-protein |
| XP_001323967.1 | TVAG_106390 | Pkeilini_DN1297_c0_g1_i1.p1 | RAB-36-and,-putative |
| XP_001323815.1 | TVAG_081640 | Pkeilini_DN46067_c0_g1_i1.p1 | conserved-hypothetical-protein |
| XP_001323810.1 | TVAG_081590 | Pkeilini_DN34893_c0_g2_i1.p1 | Rab5b,-putative |
| XP_001323776.1 | TVAG_006280 | Pkeilini_DN9662_c1_g1_i1.p1 | RAB,-putative |
| XP_001323774.1 | TVAG_006260 | Pkeilini_DN12195_c0_g3_i1.p1 | GTP-binding-protein-ypt10,-putative |
| XP_001323750.1 | TVAG_006020 | Pkeilini_DN172145_c0_g1_i1.p1 | vacuolar-ATP-synthase-subunit-ac39,-putative |
| XP_001323626.1 | TVAG_379850 | Pkeilini_DN1175_c2_g1_i1.p1 | Rab32,-putative |
| XP_001323600.1 | TVAG_379590 | Pkeilini_DN3503_c0_g1_i1.p2 | GTPase_rho,-putative |
| XP_001323596.1 | TVAG_379550 | Pkeilini_DN195601_c0_g1_i1.p1 | tyrosine-aminotransferase,-putative |

| | | | |
|---|---|---|---|
| XP_001323527.1 | TVAG_343980 | Pkeilini_DN164738_c0_g1_i1.p1 | conserved-hypothetical-protein |
| XP_001323468.1 | TVAG_127520 | Pkeilini_DN407_c0_g1_i1.p1 | N-ethylmaleimide-reductase,-putative |
| XP_001323391.1 | TVAG_498620 | Pkeilini_DN403_c0_g1_i1.p1 | ankyrin-repeat-cotaining-protein,-putative |
| XP_001323367.1 | TVAG_498380 | Pkeilini_DN66760_c1_g1_i1.p1 | Rab20,-putative |
| XP_001323255.1 | TVAG_410350 | Pkeilini_DN1857_c0_g1_i1.p2 | conserved-hypothetical-protein |
| XP_001323219.1 | TVAG_446990 | Pkeilini_DN187988_c0_g1_i1.p1 | conserved-hypothetical-protein |
| XP_001323218.1 | TVAG_446980 | Pkeilini_DN187988_c0_g1_i1.p1 | Rab9,-putative |
| XP_001323182.1 | TVAG_446610 | Pkeilini_DN2285_c0_g2_i1.p1 | small-GTPase-rabd,-putative |
| XP_001322993.1 | TVAG_121740 | Pkeilini_DN171252_c0_g1_i1.p1 | Rab5,-putative |
| XP_001322920.1 | TVAG_364210 | Pkeilini_DN14322_c0_g1_i1.p1 | Rabx37-protein,-putative |
| XP_001322907.1 | TVAG_157610 | Pkeilini_DN953_c0_g5_i1.p2 | rho4,-putative |
| XP_001322830.1 | TVAG_282070 | Pkeilini_DN497_c1_g2_i1.p1 | Rab8,-putative |
| XP_001322821.1 | TVAG_281980 | Pkeilini_DN197200_c0_g1_i1.p1 | centrosomal-protein-of-135-kDa,-putative |
| XP_001322722.1 | TVAG_484100 | Pkeilini_DN953_c0_g4_i2.p1 | Rabx26-protein,-putative |
| XP_001322593.1 | TVAG_109540 | Pkeilini_DN41445_c0_g1_i1.p1 | serine-hydroxymethyltransferase,-putative |
| XP_001322449.1 | TVAG_118780 | Pkeilini_DN108422_c0_g1_i1.p1 | calmodulin,-putative |
| XP_001322271.1 | TVAG_329350 | Pkeilini_DN7422_c1_g1_i1.p2 | Rab5,-putative |
| XP_001322074.1 | TVAG_259320 | Pkeilini_DN676_c0_g1_i2.p1 | Rabx18-protein,-putative |
| XP_001322061.1 | TVAG_259190 | Pkeilini_DN541_c0_g2_i1.p1 | succinate-thiokinase-¿-subunit |
| XP_001321827.1 | TVAG_056480 | Pkeilini_DN941_c2_g5_i1.p1 | ran,-putative |

| | | | |
|---|---|---|---|
| XP_001321627.1 | TVAG_420260 | Pkeilini_DN183133_c0_g1_i1.p1 | ATP-synthase-beta-subunit,-putative |
| XP_001321621.1 | TVAG_420200 | Pkeilini_DN608_c0_g1_i1.p1 | conserved-hypothetical-protein |
| XP_001321561.1 | TVAG_395100 | Pkeilini_DN1052_c0_g1_i1.p1 | Rab17,-putative |
| XP_001321469.1 | TVAG_133030 | Pkeilini_DN257_c0_g1_i1.p2 | NADH-ubiquinone-oxidoreductase-flavoprotein,-putative |
| XP_001321321.1 | TVAG_230580 | Pkeilini_DN688_c0_g3_i1.p1 | pyruvate-flavodoxin-oxidoreductase,-putative |
| XP_001321292.1 | TVAG_180430 | Pkeilini_DN953_c0_g2_i1.p1 | RAB-36-and,-putative |
| XP_001321289.1 | TVAG_180400 | Pkeilini_DN173495_c0_g1_i1.p1 | Phospholipase-C-precursor,-putative |
| XP_001320926.1 | TVAG_079630 | Pkeilini_DN124701_c0_g1_i1.p1 | conserved-hypothetical-protein |
| XP_001320922.1 | TVAG_079570 | Pkeilini_DN211774_c0_g1_i1.p1 | Rabx19-protein,-putative |
| XP_001320829.1 | TVAG_239660 | Pkeilini_DN176_c0_g1_i1.p1 | cysteine-desulfurylase,-putative |
| XP_001320436.1 | TVAG_308190 | Pkeilini_DN174_c0_g1_i1.p1 | ral,-putative |
| XP_001320125.1 | TVAG_064490 | Pkeilini_DN235_c0_g1_i1.p1 | rubrerythrin,-putative |
| XP_001319879.1 | TVAG_430220 | Pkeilini_DN3256_c0_g1_i1.p1 | RAB,-putative |
| XP_001319653.1 | TVAG_419720 | Pkeilini_DN171937_c0_g1_i1.p1 | aspartate-aminotransferase,-putative |
| XP_001319283.1 | TVAG_311860 | Pkeilini_DN65569_c0_g2_i1.p1 | conserved-hypothetical-protein |
| XP_001319140.1 | TVAG_057110 | Pkeilini_DN182581_c0_g1_i1.p1 | Clan-SC,-family-S33,-methylesterase-like-serine-peptidase |
| XP_001319122.1 | TVAG_056930 | Pkeilini_DN14322_c0_g1_i1.p1 | RAB-36-and,-putative |
| XP_001319084.1 | TVAG_340390 | Pkeilini_DN842_c0_g4_i1.p1 | heat-shock-protein-70-(HSP70)-4,-putative |
| XP_001319083.1 | TVAG_340380 | Pkeilini_DN91169_c0_g1_i2.p1 | conserved-hypothetical-protein |
| XP_001319074.1 | TVAG_340290 | Pkeilini_DN269_c0_g1_i2.p1 | malic-enzyme,-putative |

| | | | |
|---|---|---|---|
| XP_001318961.1 | TVAG_310250 | Pkeilini_DN111_c0_g1_i1.p1 | phosphoenolpyruvate-carboxykinase,-putative |
| XP_001318941.1 | TVAG_310050 | Pkeilini_DN13155_c3_g1_i1.p1 | nitrate,-fromate,-iron-dehydrogenase,-putative |
| XP_001318893.1 | TVAG_211200 | Pkeilini_DN953_c0_g8_i1.p1 | Rabx21-protein,-putative |
| XP_001318848.1 | TVAG_056190 | Pkeilini_DN725_c1_g1_i1.p1 | Clan-MH,-family-M20,-peptidase-T-like-metallopeptidase |
| XP_001318715.1 | TVAG_257310 | Pkeilini_DN941_c2_g3_i1.p1 | Rab21,-putative |
| XP_001318541.1 | TVAG_065750 | Pkeilini_DN104585_c0_g1_i1.p1 | sucrose-transport-protein,-putative |
| XP_001318394.1 | TVAG_098820 | Pkeilini_DN195601_c0_g1_i1.p1 | tyrosine-aminotransferase,-putative |
| XP_001318376.1 | TVAG_068130 | Pkeilini_DN269_c0_g1_i2.p1 | malic-enzyme,-putative |
| XP_001317955.1 | TVAG_100550 | Pkeilini_DN144216_c0_g1_i1.p1 | glycine-cleavage-system-H-protein,-putative |
| XP_001317694.1 | TVAG_286490 | Pkeilini_DN3462_c4_g1_i1.p1 | RAB,-putative |
| XP_001317473.1 | TVAG_019960 | Pkeilini_DN317_c0_g2_i3.p1 | conserved-hypothetical-protein |
| XP_001317292.1 | TVAG_191660 | Pkeilini_DN184795_c0_g1_i1.p1 | groes-chaperonin,-putative |
| XP_001317232.1 | TVAG_416520 | Pkeilini_DN1175_c2_g1_i1.p1 | RAB-18,-putative |
| XP_001317174.1 | TVAG_040030 | Pkeilini_DN1216_c0_g1_i1.p1 | Iron-sulfur-flavoprotein |
| XP_001317169.1 | TVAG_039980 | Pkeilini_DN108_c0_g2_i1.p3 | superoxide-dismutase,-putative |
| XP_001317041.1 | TVAG_226310 | Pkeilini_DN474_c2_g1_i1.p1 | conserved-hypothetical-protein |
| XP_001316822.1 | TVAG_233350 | Pkeilini_DN50982_c0_g1_i1.p1 | Clan-ME,-family-M16,-insulinase-like-metallopeptidase |
| XP_001316713.1 | TVAG_453070 | Pkeilini_DN258_c0_g1_i1.p1 | Rab2,-putative |
| XP_001316606.1 | TVAG_350580 | Pkeilini_DN941_c0_g1_i1.p1 | Rab21,-putative |
| XP_001316281.1 | TVAG_203620 | Pkeilini_DN392_c0_g1_i1.p1 | rubisco-subunit-binding-protein-alpha-subunit,-ruba,-putative |

| | | | |
|---|---|---|---|
| XP_001316041.1 | TVAG_454570 | Pkeilini_DN1322_c0_g1_i1.p1 | Embryonic-protein-DC-8,-putative |
| XP_001315530.1 | TVAG_193770 | Pkeilini_DN953_c0_g5_i1.p2 | RAB,-putative |
| XP_001315422.1 | TVAG_049830 | Pkeilini_DN266_c0_g1_i1.p1 | disulfide-oxidoreductase,-putative |
| XP_001315408.1 | TVAG_049690 | Pkeilini_DN217534_c0_g1_i1.p1 | thiamin-pyrophosphokinase,-putative |
| XP_001315345.1 | TVAG_328940 | Pkeilini_DN160822_c0_g1_i1.p1 | alcohol-dehydrogenase,-putative |
| XP_001315025.1 | TVAG_008100 | Pkeilini_DN1297_c0_g1_i1.p1 | Rabx38-protein,-putative |
| XP_001314995.1 | TVAG_370000 | Pkeilini_DN12195_c0_g1_i1.p1 | Rabx30-protein,-putative |
| XP_001314846.1 | TVAG_260830 | Pkeilini_DN5872_c0_g2_i1.p1 | conserved-hypothetical-protein |
| XP_001314811.1 | TVAG_254890 | Pkeilini_DN688_c0_g3_i1.p1 | pyruvate-flavodoxin-oxidoreductase,-putative |
| XP_001314747.1 | TVAG_020610 | Pkeilini_DN1975_c0_g1_i1.p2 | RAB-18,-putative |
| XP_001314705.1 | TVAG_055550 | Pkeilini_DN3462_c0_g1_i1.p1 | Rab8,-putative |
| XP_001314029.1 | TVAG_412220 | Pkeilini_DN269_c0_g1_i2.p1 | malic-enzyme,-putative |
| XP_001313967.1 | TVAG_442270 | Pkeilini_DN4169_c0_g1_i1.p1 | GTP-binding-protein-yptv3,-putative |
| XP_001313958.1 | TVAG_442170 | Pkeilini_DN38635_c0_g1_i1.p1 | conserved-hypothetical-protein |
| XP_001313802.1 | TVAG_265760 | Pkeilini_DN407_c0_g1_i1.p1 | N-ethylmaleimide-reductase,-putative |
| XP_001313682.1 | TVAG_096630 | Pkeilini_DN217534_c0_g1_i1.p1 | thiamin-pyrophosphokinase,-putative |
| XP_001313584.1 | TVAG_208470 | Pkeilini_DN160426_c0_g1_i1.p1 | threonyl-tRNA-synthetase,-putative |
| XP_001313551.1 | TVAG_181000 | Pkeilini_DN206_c0_g5_i1.p1 | RAB-19,-41-and,-putative |
| XP_001313356.1 | TVAG_114310 | Pkeilini_DN4486_c0_g1_i1.p1 | peroxiredoxins,-prx-1,-prx-2,-prx-3,-putative |
| XP_001313153.1 | TVAG_257780 | Pkeilini_DN272_c2_g3_i1.p1 | biotin-synthase,-putative |

| | | | |
|---|---|---|---|
| XP_001312991.1 | TVAG_196220 | Pkeilini_DN207877_c0_g1_i1.p1 | protein-brittle-1,-chloroplast-precursor,-putative |
| XP_001312927.1 | TVAG_029020 | Pkeilini_DN1080_c0_g1_i1.p1 | small-GTPase-rabh,-putative |
| XP_001312785.1 | TVAG_424920 | Pkeilini_DN66760_c1_g1_i1.p1 | RAB-3-and,-putative |
| XP_001312753.1 | TVAG_088050 | Pkeilini_DN392_c0_g1_i1.p1 | chaperonin,-putative |
| XP_001312620.1 | TVAG_468600 | Pkeilini_DN182200_c0_g1_i1.p1 | WD-repeat-protein,-putative |
| XP_001312469.1 | TVAG_241150 | Pkeilini_DN26_c1_g1_i1.p2 | Rabx24-protein,-putative |
| XP_001312254.1 | TVAG_152430 | Pkeilini_DN159673_c0_g1_i1.p1 | lysyl-tRNA-synthetase,-putative |
| XP_001312198.1 | TVAG_112840 | Pkeilini_DN187988_c0_g1_i1.p1 | Rabx15-protein,-putative |
| XP_001312168.1 | TVAG_296220 | Pkeilini_DN160340_c0_g1_i1.p1 | NADH-dehydrogenase-24-kDa-subunit,-putative |
| XP_001312142.1 | TVAG_236570 | Pkeilini_DN4000_c0_g1_i1.p1 | Rab32,-putative |
| XP_001312073.1 | TVAG_115470 | Pkeilini_DN195996_c0_g1_i1.p1 | conserved-hypothetical-protein |
| XP_001311960.1 | TVAG_008790 | Pkeilini_DN71584_c0_g1_i1.p1 | conserved-hypothetical-protein |
| XP_001311873.1 | TVAG_321030 | Pkeilini_DN195918_c0_g1_i1.p1 | conserved-hypothetical-protein |
| XP_001311871.1 | TVAG_321010 | Pkeilini_DN49090_c0_g2_i1.p1 | AMP-dependent-ligase/synthetase,-putative |
| XP_001311861.1 | TVAG_242960 | Pkeilini_DN688_c0_g3_i1.p1 | pyruvate-flavodoxin-oxidoreductase,-putative |
| XP_001311772.1 | TVAG_409800 | Pkeilini_DN207636_c0_g1_i1.p1 | RAB,-putative |
| XP_001311734.1 | TVAG_479760 | Pkeilini_DN47132_c0_g1_i1.p1 | conserved-hypothetical-protein |
| XP_001311148.1 | TVAG_263350 | Pkeilini_DN3153_c0_g1_i1.p1 | conserved-hypothetical-protein |
| XP_001311109.1 | TVAG_047800 | Pkeilini_DN2125_c0_g1_i1.p1 | Rab11,-putative |
| XP_001310420.1 | TVAG_335500 | Pkeilini_DN17664_c0_g1_i1.p1 | conserved-hypothetical-protein |
| XP_001310180.1 | TVAG_037570 | Pkeilini_DN13155_c0_g4_i1.p1 | NADH-ubiquinone-oxidoreductase,-putative |

| | | | |
|---|---|---|---|
| XP_001310176.1 | TVAG_037530 | Pkeilini_DN8689_c0_g1_i1.p1 | conserved-hypothetical-protein |
| XP_001309934.1 | TVAG_440200 | Pkeilini_DN164467_c0_g1_i1.p1 | conserved-hypothetical-protein |
| XP_001309798.1 | TVAG_390750 | Pkeilini_DN26408_c0_g3_i1.p2 | RAB-18,-putative |
| XP_001309776.1 | TVAG_470110 | Pkeilini_DN197003_c0_g1_i1.p1 | conserved-hypothetical-protein |
| XP_001309717.1 | TVAG_216900 | Pkeilini_DN953_c0_g9_i1.p1 | ras,-putative |
| XP_001309656.1 | TVAG_075320 | Pkeilini_DN3946_c0_g1_i1.p2 | vacuolar-proton-ATPase,-putative |
| XP_001309521.1 | TVAG_018050 | Pkeilini_DN217915_c0_g1_i1.p2 | RAB-18,-putative |
| XP_001309408.1 | TVAG_277590 | Pkeilini_DN160251_c0_g1_i1.p1 | cation-transporting-ATPase,-putative |
| XP_001309398.1 | TVAG_024790 | Pkeilini_DN494_c0_g1_i1.p2 | septum-promoting-GTP-binding-protein,-putative |
| XP_001309295.1 | TVAG_337970 | Pkeilini_DN106_c0_g1_i1.p1 | conserved-hypothetical-protein |
| XP_001309218.1 | TVAG_264120 | Pkeilini_DN34504_c0_g3_i1.p1 | pecanex,-putative |
| XP_001309182.1 | TVAG_205390 | Pkeilini_DN350_c0_g1_i2.p1 | GTPase-mss1/trme,-putative |
| XP_001308879.1 | TVAG_077910 | Pkeilini_DN4238_c0_g1_i1.p2 | conserved-hypothetical-protein |
| XP_001308636.1 | TVAG_256470 | Pkeilini_DN217534_c0_g1_i1.p1 | thiamin-pyrophosphokinase,-putative |
| XP_001308553.1 | TVAG_047210 | Pkeilini_DN2207_c0_g1_i1.p1 | 2-hydroxyacid-dehydrogenase,-putative |
| XP_001308528.1 | TVAG_459470 | Pkeilini_DN1975_c0_g1_i1.p2 | Rab9,-putative |
| XP_001308237.1 | TVAG_032090 | Pkeilini_DN622_c0_g1_i1.p1 | Co-chaperone-protein-HscB-Hsc20,-mitochondrial-precursor,-putative |
| XP_001308201.1 | TVAG_051830 | Pkeilini_DN3462_c4_g1_i1.p1 | RAC,-putative |
| XP_001308200.1 | TVAG_051820 | Pkeilini_DN172140_c0_g1_i1.p1 | tricarboxylate-transport-protein,-putative |
| XP_001308167.1 | TVAG_336320 | Pkeilini_DN3171_c0_g2_i1.p1 | Hydroxylamine-reductase,-putative |

| | | | |
|---|---|---|---|
| XP_001307912.1 | TVAG_048600 | Pkeilini_DN1721_c0_g1_i1.p1 | small-GTPase-rabi,-putative |
| XP_001307775.1 | TVAG_049140 | Pkeilini_DN108_c0_g2_i1.p3 | superoxide-dismutase-[fe],-putative |
| XP_001307690.1 | TVAG_346230 | Pkeilini_DN41178_c0_g1_i1.p2 | conserved-hypothetical-protein |
| XP_001307488.1 | TVAG_300910 | Pkeilini_DN199171_c0_g1_i1.p1 | Rabx18-protein,-putative |
| XP_001307405.1 | TVAG_458060 | Pkeilini_DN157931_c0_g2_i1.p1 | conserved-hypothetical-protein |
| XP_001307320.1 | TVAG_194760 | Pkeilini_DN64940_c0_g1_i1.p1 | guanine-nucleotide-exchange-factor,-putative |
| XP_001307251.1 | TVAG_466790 | Pkeilini_DN206198_c0_g1_i1.p1 | pyruvate-flavodoxin-oxidoreductase,-putative |
| XP_001307088.1 | TVAG_105770 | Pkeilini_DN206198_c0_g1_i1.p1 | pyruvate-flavodoxin-oxidoreductase,-putative |
| XP_001307064.1 | TVAG_080400 | Pkeilini_DN953_c0_g10_i1.p1 | Rabx31-protein,-putative |
| XP_001306984.1 | TVAG_151010 | Pkeilini_DN2353_c0_g1_i3.p3 | Rabx19-protein,-putative |
| XP_001306776.1 | TVAG_224980 | Pkeilini_DN160002_c0_g1_i1.p1 | Clan-MH,-family-M20,-peptidase-T-like-metallopeptidase |
| XP_001306669.1 | TVAG_455090 | Pkeilini_DN4238_c0_g1_i1.p2 | conserved-hypothetical-protein |
| XP_001306447.1 | TVAG_132350 | Pkeilini_DN830_c0_g1_i1.p1 | conserved-hypothetical-protein |
| XP_001306356.1 | TVAG_104710 | Pkeilini_DN1085_c0_g1_i1.p1 | Rabx3-protein,-putative |
| XP_001306230.1 | TVAG_277050 | Pkeilini_DN27394_c0_g1_i1.p1 | Citrate-lyase-beta-chain,-putative |
| XP_001305871.1 | TVAG_489800 | Pkeilini_DN222480_c0_g1_i1.p1 | NADH-dehydrogenase-51-kDa-subunit,-putative |
| XP_001305709.1 | TVAG_361590 | Pkeilini_DN13155_c0_g4_i1.p1 | NADH-ubiquinone-oxidoreductase,-putative |
| XP_001305704.1 | TVAG_361540 | Pkeilini_DN419_c0_g1_i1.p2 | iron-sulfur-assembly-protein,-putative |
| XP_001305570.1 | TVAG_349870 | Pkeilini_DN26_c0_g1_i2.p1 | Rabx15-protein,-putative |
| XP_001305426.1 | TVAG_344520 | Pkeilini_DN914_c0_g1_i1.p1 | conserved-hypothetical-protein |

| | | | |
|---|---|---|---|
| XP_001305403.1 | TVAG_331490 | Pkeilini_DN9662_c1_g1_i1.p1 | Rabx18-protein,-putative |
| XP_001305368.1 | TVAG_327470 | Pkeilini_DN771_c0_g1_i1.p1 | alcohol-dehydrogenase,-putative |
| XP_001305213.1 | TVAG_262750 | Pkeilini_DN160677_c0_g1_i1.p1 | vacuolar-ATP-synthase-subunit-H,-putative |
| XP_001305174.1 | TVAG_164890 | Pkeilini_DN182694_c0_g1_i1.p1 | Acetyl-CoA-hydrolase,-putative |
| XP_001305114.1 | TVAG_386080 | Pkeilini_DN170837_c0_g1_i1.p1 | Clan-MG,-familly-M24,-aminopeptidase-P-like-metallopeptidase |
| XP_001305092.1 | TVAG_383350 | Pkeilini_DN196009_c0_g1_i1.p1 | RAB-2,4,14,-putative |
| XP_001304655.1 | TVAG_384490 | Pkeilini_DN429_c0_g1_i1.p1 | Rab9,-putative |
| XP_001304618.1 | TVAG_092740 | Pkeilini_DN219560_c0_g1_i1.p1 | small-GTPase-rabi,-putative |
| XP_001304246.1 | TVAG_065320 | Pkeilini_DN2353_c0_g1_i3.p3 | Rab21,-putative |
| XP_001304133.1 | TVAG_220970 | Pkeilini_DN16958_c0_g1_i1.p2 | RAB,-putative |
| XP_001304067.1 | TVAG_147840 | Pkeilini_DN210433_c0_g1_i1.p1 | Rab21,-putative |
| XP_001304062.1 | TVAG_147790 | Pkeilini_DN6516_c0_g2_i1.p1 | cysteine/methionine-metabolism-pyridoxal-5-phosphate-enzymes,-putative |
| XP_001303981.1 | TVAG_144730 | Pkeilini_DN541_c0_g1_i1.p1 | succinate-thiokinase-¿-subunit |
| XP_001303783.1 | TVAG_445730 | Pkeilini_DN199807_c0_g1_i1.p1 | groes-chaperonin,-putative |
| XP_001303674.1 | TVAG_092170 | Pkeilini_DN1127_c0_g1_i1.p1 | preprotein-translocase-secy-subunit,-putative |
| XP_001303658.1 | TVAG_134510 | Pkeilini_DN219911_c0_g1_i1.p1 | RAB-18,-putative |
| XP_001303641.1 | TVAG_328110 | Pkeilini_DN26_c0_g1_i2.p1 | RAB-18,-putative |
| XP_001303632.1 | TVAG_152690 | Pkeilini_DN407_c0_g1_i1.p1 | N-ethylmaleimide-reductase,-putative |
| XP_001303268.1 | TVAG_385350 | Pkeilini_DN183936_c0_g1_i1.p1 | thioredoxin,-putative |
| XP_001303252.1 | TVAG_234150 | Pkeilini_DN109943_c0_g1_i1.p1 | conserved-hypothetical-protein |

| | | | |
|---|---|---|---|
| XP_001303146.1 | TVAG_415960 | Pkeilini_DN3256_c0_g1_i1.p1 | Rab9,-putative |
| XP_001303059.1 | TVAG_088220 | Pkeilini_DN195601_c0_g1_i1.p1 | aspartate-aminotransferase,-putative |
| XP_001302997.1 | TVAG_371280 | Pkeilini_DN12181_c0_g1_i1.p1 | Rab2,-putative |
| XP_001302917.1 | TVAG_124590 | Pkeilini_DN4394_c0_g1_i1.p1 | Rab6,-putative |
| XP_001302913.1 | TVAG_124540 | Pkeilini_DN158973_c0_g1_i1.p1 | GTPase_rho,-putative |
| XP_001302832.1 | TVAG_320300 | Pkeilini_DN184375_c0_g1_i1.p1 | GTP-binding-protein-Rab2,-putative |
| XP_001302560.1 | TVAG_386000 | Pkeilini_DN159371_c0_g1_i1.p1 | Receptor-expression-enhancing-protein,-putative |
| XP_001302496.1 | TVAG_356810 | Pkeilini_DN144134_c0_g2_i1.p1 | NimA-like-protein |
| XP_001302364.1 | TVAG_327760 | Pkeilini_DN1216_c0_g1_i1.p1 | Iron-sulfur-flavoprotein |
| XP_001302013.1 | TVAG_261280 | Pkeilini_DN2117_c0_g1_i1.p1 | GTP-binding-protein-ypt11,-putative |
| XP_001301760.1 | TVAG_025980 | Pkeilini_DN11926_c1_g1_i2.p1 | glutamate-dehydrogenase,-putative |
| XP_001301244.1 | TVAG_158270 | Pkeilini_DN222101_c0_g1_i1.p1 | RAB,-putative |
| XP_001301100.1 | TVAG_041340 | Pkeilini_DN199807_c0_g1_i1.p1 | groes-chaperonin,-putative |
| XP_001301044.1 | TVAG_422690 | Pkeilini_DN1052_c0_g1_i1.p1 | Rab9,-putative |
| XP_001301038.1 | TVAG_422630 | Pkeilini_DN622_c0_g1_i1.p1 | Co-chaperone-protein-HscB-Hsc20,-mitochondrial-precursor,-putative |
| XP_001300804.1 | TVAG_293370 | Pkeilini_DN207058_c0_g1_i1.p1 | nucleoside-diphosphate-kinase,-putative |
| XP_001300601.1 | TVAG_001130 | Pkeilini_DN53254_c0_g1_i1.p1 | conserved-hypothetical-protein |
| XP_001300482.1 | TVAG_318670 | Pkeilini_DN626_c0_g1_i1.p1 | succinate-thiokinase-a-subunit |
| XP_001300294.1 | TVAG_015270 | Pkeilini_DN6278_c0_g1_i2.p2 | small-GTPase-rabh,-putative |
| XP_001300248.1 | TVAG_169740 | Pkeilini_DN219911_c0_g1_i1.p1 | RAB-19,-41-and,-putative |

| | | | |
|---|---|---|---|
| XP_001299687.1 | TVAG_440690 | Pkeilini_DN26_c4_g1_i1.p1 | small-GTPase-rabh,-putative |
| XP_001299604.1 | TVAG_527180 | Pkeilini_DN327_c2_g1_i1.p1 | RAB,-putative |
| XP_001299513.1 | TVAG_177600 | Pkeilini_DN183618_c0_g1_i1.p1 | glycine-cleavage-system-H-protein,-putative |
| XP_001299483.1 | TVAG_450220 | Pkeilini_DN96108_c0_g1_i1.p1 | conserved-hypothetical-protein |
| XP_001299482.1 | TVAG_399860 | Pkeilini_DN71456_c0_g1_i1.p1 | Ferredoxin-2 |
| XP_001299372.1 | TVAG_499340 | Pkeilini_DN497_c1_g1_i1.p1 | hypothetical-protein |
| XP_001299219.1 | TVAG_528800 | Pkeilini_DN3462_c4_g1_i1.p1 | RAB-2,4,14,-putative |
| XP_001299204.1 | TVAG_450060 | Pkeilini_DN13377_c0_g1_i1.p1 | conserved-hypothetical-protein |
| XP_001298987.1 | TVAG_126970 | Pkeilini_DN207052_c0_g1_i1.p1 | RAB-GDP-dissociation-inhibitor,-putative |
| XP_001298262.1 | TVAG_022530 | Pkeilini_DN30686_c0_g1_i1.p1 | conserved-hypothetical-protein |
| XP_001297863.1 | TVAG_377380 | Pkeilini_DN1554_c0_g1_i1.p1 | conserved-hypothetical-protein |
| XP_001297704.1 | TVAG_085320 | Pkeilini_DN2659_c0_g1_i1.p1 | small-GTPase-rabi,-putative |
| XP_001297366.1 | TVAG_530140 | Pkeilini_DN160991_c0_g1_i1.p1 | conserved-hypothetical-protein |
| XP_001297292.1 | TVAG_504530 | Pkeilini_DN211774_c0_g1_i1.p1 | Rabx26-protein,-putative |
| XP_001296818.1 | TVAG_060820 | Pkeilini_DN1085_c1_g1_i1.p1 | RAB,-putative |
| XP_001296212.1 | TVAG_082020 | Pkeilini_DN159740_c0_g1_i1.p1 | conserved-hypothetical-protein |
| XP_001294582.1 | TVAG_547520 | Pkeilini_DN218014_c0_g1_i1.p1 | threonine-synthase,-putative |
| XP_001294517.1 | TVAG_416100 | Pkeilini_DN568_c0_g1_i1.p1 | malic-enzyme,-putative |

**Table S4: List of annotated genes gained in the trichomonas lineage after the split from their common ancestor with *P. keilini***

| Gene family | Annotation | *T. vaginalis* gene ID | KEGG_ko |
|---|---|---|---|
| OG_792 | Acetyltransf_1 | TvagiEBg036290 | ko:K20793 |
| OG_5491 | Acetyltransf_1,Pkinase,TPR_8 | TvagiEBg053982 | ko:K04345 |
| OG_24374 | biological adhesion | TvagiEBg038277 | NA |
| OG_24315 | biological adhesion | TvagiEBg036641 | ko:K10402 |
| OG_6750 | biological adhesion | TvagiEBg023227 | NA |
| OG_7081 | biological adhesion | TvagiEBg023531 | NA |
| OG_7139 | biological adhesion | TvagiEBg055182 | ko:K14000 |
| OG_7120 | biological adhesion | TvagiEBg043549 | NA |
| OG_16843 | biological adhesion | TvagiEBg047580 | NA |
| OG_16996 | biological adhesion | TvagiEBg036104 | ko:K14000 |
| OG_17311 | biological adhesion | TvagiEBg041328 | ko:K15152 |
| OG_170 | cAMP-dependent protein kinase activity | TvagiEBg039290 | ko:K04345,ko:K10409,ko:K19584 |
| OG_1811 | Chromo,DUF4208,Helicase_C,PHD,SNF2_N | TvagiEBg028330 | ko:K11367 |
| OG_2044 | DNA polymerase type B, organellar and viral | TvagiEBg047836 | NA |
| OG_3706 | twin BRCT domain | TvagiEBg028318 | ko:K10728 |

| | | | |
|---|---|---|---|
| OG_2184 | endoplasmic reticulum-plasma membrane tethering | TvagiEBg028132 | ko:K12486,ko:K19938 |
| OG_3706 | twin BRCT domain | TvagiEBg028318 | ko:K10728 |
| OG_1726 | spectrin binding | TvagiEBg042842,TvagiEBg055069 | ko:K15502,ko:K15503 |
| OG_6482 | Glycine rich protein | TvagiEBg030396 | NA |
| OG_6775 | GTPase activity | TvagiEBg034972 | NA |
| OG_6405 | GTPase activity | TvagiEBg041930 | NA |
| OG_2058 | islet amyloid polypeptide processing | TvagiEBg048541,TvagiEBg048542 | ko:K01341,ko:K01360,ko:K08673 |
| OG_1775 | Exhibits S-adenosyl-L-methionine-dependent methyltransferase activity | TvagiEBg025243 | NA |
| OG_2007 | MyD88-dependent toll-like receptor signaling pathway | TvagiEBg044672,TvagiEBg044653 | ko:K18809 |
| OG_5666 | Myb-like DNA-binding domain | TvagiEBg022475,TvagiEBg041712,TvagiEBg055335,TvagiEBg046332 | ko:K09422 |
| OG_16767 | nerve growth factor signaling pathway | TvagiEBg056182 | ko:K15503,ko:K21440 |
| OG_6132 | nuclear import signal receptor activity | TvagiEBg055732 | NA |
| OG_24385 | PDZ domain binding | TvagiEBg057545 | ko:K16072,ko:K19878,ko:K20478 |
| OG_1974 | Phage tail repeat like | TvagiEBg019118 | NA |

| | | | |
|---|---|---|---|
| OG_24226 | spectrin binding | TvagiEBg046228 | NA |
| OG_5955 | protein kinase activity | TvagiEBg050562,TvagiEBg050583 | ko:K04345 |
| OG_5921 | protein serine/threonine kinase activity | TvagiEBg025205,TvagiEBg054198,TvagiEBg046872 | NA |
| OG_1756 | protein serine/threonine kinase activity | TvagiEBg023811,TvagiEBg041668 | ko:K02216 |
| OG_1026 | protein serine/threonine kinase activity | TvagiEBg028031,TvagiEBg049667 | ko:K08813,ko:K16312 |
| OG_4577 | protein serine/threonine kinase activity | TvagiEBg038832,TvagiEBg048970 | ko:K04428,ko:K17533,ko:K08798 |
| OG_6690 | protein serine/threonine kinase activity | TvagiEBg056404 | ko:K08857,ko:K20879 |
| OG_24246 | protein ubiquitination | TvagiEBg000145 | NA |
| OG_24247 | protein ubiquitination | TvagiEBg033567,TvagiEBg035052 | ko:K20129,ko:K15502,ko:K15503 |
| OG_1735 | protein ubiquitination | TvagiEBg021138,TvagiEBg024708,TvagiEBg012479,TvagiEBg001122,TvagiEBg001731,TvagiEBg006733,TvagiEBg000512,TvagiEBg008998,TvagiEBg002326 | ko:K10325,ko:K12591 |
| OG_6128 | protein ubiquitination | TvagiEBg049297,TvagiEBg043655 | NA |
| OG_16927 | protein ubiquitination | TvagiEBg039035 | ko:K15502,ko:K15503,ko:K20032,ko:K21440 |
| OG_2368 | protein serine/threonine kinase activity | TvagiEBg032765,TvagiEBg047551,TvagiEBg016996,TvagiEBg058648,TvagiEBg059205,TvagiEBg052716 | ko:K04345,ko:K07376,ko:K13302,ko:K13303 |
| OG_7079 | regulation of centriole replication | TvagiEBg021289 | ko:K06631,ko:K07298,ko:K08269,ko:K08850,ko:K13412,ko:K21358 |

| | | | |
|---|---|---|---|
| OG_6445 | regulation of choline O-acetyltransferase activity | TvagiEBg049095,TvagiEBg050875 | ko:K01404,ko:K08654,ko:K12813 |
| OG_4922 | Reprolysin_4 | TvagiEBg044791,TvagiEBg044795,TvagiEBg050876 | NA |
| OG_19677 | Reversible hydration of carbon dioxide | TvagiEBg041073,TvagiEBg054911 | ko:K01673 |
| OG_5952 | Ribonuclease H protein | TvagiEBg028675,TvagiEBg029501,TvagiEBg028461,TvagiEBg038876,TvagiEBg037831,TvagiEBg030542,TvagiEBg040960,TvagiEBg032543,TvagiEBg027125 | ko:K12879 |
| OG_18737 | Ribosomal L29e protein family rpl29 | TvagiEBg037733,TvagiEBg034570.TvagiEBg047046 | ko:K02905 |
| OG_1071 | Right handed beta helix region | TvagiEBg032352,TvagiEBg045961,TvagiEBg045962,TvagiEBg020968,TvagiEBg028656,TvagiEBg002344,TvagiEBg017538,TvagiEBg051902,TvagiEBg052658,TvagiEBg055958 | NA |
| OG_19828 | RNA polymerase I core binding | TvagiEBg026015 | ko:K11294 |
| OG_24227 | RNA polymerase II transcription regulator recruiting activity | TvagiEBg000087,TvagiEBg054872,TvagiEBg058883 | NA |
| OG_6053 | RNA binding | TvagiEBg023316 | ko:K13154 |
| OG_19443 | sensory perception of sound | TvagiEBg028625,TvagiEBg051265 | ko:K05747 |
| OG_1990 | serine-type endopeptidase activity | TvagiEBg052350,TvagiEBg052355,TvagiEBg052562,TvagiEBg052386,TvagiEBg052378 | NA |
| OG_24486 | spectrin binding | TvagiEBg054770 | NA |

| | | | |
|---|---|---|---|
| OG_1846 | spectrin binding | TvagiEBg037450,TvagiEBg037444,TvagiEBg037452,TvagiEBg030906,TvagiEBg037447,TvagiEBg059166,TvagiEBg032170,TvagiEBg028616,TvagiEBg027013 | ko:K15502,ko:K15503 |
| OG_124 | spectrin binding,protein ubiquitination | TvagiEBg036895,TvagiEBg039181,TvagiEBg056857,TvagiEBg058173,TvagiEBg051128,TvagiEBg025287,TvagiEBg039180,TvagiEBg025285,TvagiEBg025286,TvagiEBg046793,TvagiEBg045177,TvagiEBg055607,TvagiEBg059132,TvagiEBg049319 | ko:K15502,ko:K15503,ko:K20032,ko:K21440 |
| OG_1726 | spectrin binding | TvagiEBg042842,TvagiEBg055069 | ko:K15502,ko:K15503 |
| OG_6626 | spectrin binding | TvagiEBg031656 | NA |
| OG_7116 | spectrin binding | TvagiEBg041913 | NA |
| OG_17052 | spectrin binding | TvagiEBg045359 | ko:K10380 |
| OG_2826 | Sulfatase | TvagiEBg043468,TvagiEBg056542,TvagiEBg038218,TvagiEBg049604,TvagiEBg020034,TvagiEBg037070,TvagiEBg040806,TvagiEBg056543,TvagiEBg039040,TvagiEBg044578,TvagiEBg019289,TvagiEBg044686,TvagiEBg051277,TvagiEBg050423,TvagiEBg040807,TvagiEBg049450,TvagiEBg015897,TvagiEBg021826,TvagiEBg019340 | NA |
| OG_5283 | thiosulfate sulfurtransferase activity | TvagiEBg056003 | ko:K01011 |
| OG_701 | thiomorpholine-carboxylate dehydrogenase activity | TvagiEBg027824,TvagiEBg042806 | ko:K01750,ko:K18258 |

| | | | |
|---|---|---|---|
| OG_218 | trehalase (brush-border membrane glycoprotein) | TvagiEBg044074,TvagiEBg044075 | ko:K01194 |
| OG_2106 | unfolded protein binding, ATP binding | TvagiEBg052412,TvagiEBg036462,TvagiEBg045708,TvagiEBg056047,TvagiEBg055070,TvagiEBg023823,TvagiEBg038251,TvagiEBg052522,TvagiEBg022564,TvagiEBg052488,TvagiEBg052411 | ko:K04043 |

### *P. keilini* hydrogenosomal preprotein import system is similar to other hydrogenosomes

Import into the hydrogenosome requires transport across two plasma membranes, in canonical mitochondrial organisms the inner and outer membranes have evolutionarily unrelated translocase complexes, which contain a core translocase and accessory components which assist in preprotein import. The Translocase of the Outer Membrane (TOM) complex has an essential conserved translocase, TOM40, and orthologs have been functionally characterised as the preprotein translocases in isolated hydrogenosomes (Makki et al., 2019). Characteristic of these proteins is a beta-barrel fold of the pfam hmm family Porin_3, the number of paralogs of this protein varies from lineage to lineage in excavates with kinetoplastids such a *T. brucei* having two copies of a highly diverged protein termed ATOM (Pusnik et al., 2011), and *T. vaginalis* as many as six (Rada et al., 2011). We identified two partial sequences in *P. keilini* (Pfam Porin_3, E-value 1.1x10^6, 2.3x10^5). Whilst neither sequence is complete, both have beta barrel topology similar to translocases identified in *T. vaginalis* by PRED_TMBB (Bagos et al., 2004). In most eukaryotes the TOM complex has accessory proteins which assist in preprotein import and binding, these subunits seem to have independently emerged in different eukaryotic lineages though are assumedly functionally similar. Several accessory proteins have been identified to the *T. brucei* TOM complex (Schneider, 2018), though appear absent in the *T. vaginalis* (Makki et al., 2019). No strong homologs to the accessory proteins from the yeast system are present in *P. keilini* nor ATOM11, 12, 14, 46, 69 of the *T. brucei* TOM complex.

Preproteins destined for insertion into the outer membrane are handled subsequent to Tom40 import by another bacterially evolved beta barrel protein SAM50 which is the core translocase of the SAM complex (Kozjak et al., 2003). We did not detect SAM50 in *P. keilini.*

In contrast to the ancestrally bacterial beta barrel translocases of the outer membrane, import through and insertion into the inner membrane is facilitated by proteins of eukaryotic innovation termed Translocase of the Inner Membrane (TIM) complexes. In many eukaryotes the inner membrane translocases have functionally diverged to do slightly different tasks, in yeast three related proteins Tim17, 22, 23 form the core translocases to two different complexes the Tim22 complex mediating the insertion of membrane proteins and Tim23 for lumen destined proteins. In *P. keilini* two proteins were identified with homology to the inner membrane TIM 17 family (Pfam Tim17, E-values 3.3x10^8, 4.6x10^8), this compares to the 4 identified in *T. vaginalis* where it is not yet clear whether they form functionally discrete complexes (Rada et al., 2011). Like the outer membrane, the translocase complexes of the inner membrane have accessory subunits which assist in preprotein import, the most significant of these the Presequence translocase-Associated Motor (PAM) which involves both matrix and membrane associated proteins. The membrane associating components, Tim44 Pam16 (Pfam Pam16, E-value 8.4x10^5), and Pam18 were all found in *P. keilini* as well as the matrix proteins Mge1 and mtHSP70. The PAM motor has also been characterised in *T. vaginalis* (Rada et al., 2011) and is similarly complete. Some imported preproteins undergo signal sequence cleavage with a specific Mitochondrial Processing Protease (MPP). In many eukaryotes the MPP is composed of evolutionarily related $\alpha/\beta$ subunits, but other organisms including *T. vaginalis* have undergone streamlining to a single subunit (Šmíd et al., 2008). In our dataset we find only a single MPP type protease, suggesting that the reductive evolution of this complex occurred before the speciation of *P. keilini*.

In summary the *P. keilini* hydrogenosome retains the core translocases of the inner and outer membranes, and closely resembles the architecture of the *Trichomonas* hydrogenosome, with complete PAM and a single MPP, but without other accessory proteins. The copy number of the core translocases is lower than that of *T. vaginalis*. Particularly interesting in this respect are the two Tim17 family proteins, which would suggest that the scope for functional diversity of the TIM complexes is limited to those two proteins and could be another example of a reductive evolution in the preprotein import system.
